# Epigenomic translocation of H3K4me3 broad domains over oncogenes following hijacking of super-enhancers

**DOI:** 10.1101/2020.02.12.938563

**Authors:** Aneta Mikulasova, Daniel Kent, Marco Trevisan-Herraz, Nefeli Karataraki, Kent T.M. Fung, Cody Ashby, Agata Cieslak, Shmuel Yaccoby, Frits van Rhee, Maurizio Zangari, Sharmilan Thanendrarajan, Carolina Schinke, Gareth J. Morgan, Vahid Asnafi, Salvatore Spicuglia, Chris A. Brackley, Anne E. Corcoran, Sophie Hambleton, Brian A. Walker, Daniel Rico, Lisa J. Russell

## Abstract

Chromosomal translocations are important drivers of haematological malignancies whereby proto-oncogenes are activated by juxtaposition with enhancers, often called *enhancer hijacking*. We analysed the epigenomic consequences of rearrangements between the super-enhancers of the immunoglobulin heavy locus (*IGH)* and proto-oncogene *CCND1* that are common in B-cell malignancies. By integrating BLUEPRINT epigenomic data with DNA breakpoint detection, we characterised the normal chromatin landscape of the human *IGH* locus and its dynamics after pathological genomic rearrangement. We detected an H3K4me3 broad domain (BD) within the *IGH* locus of healthy B cells that was absent in samples with *IGH-CCND1* translocations. The appearance of H3K4me3-BD over *CCND1* in the latter was associated with overexpression and extensive chromatin accessibility of its gene body. We observed similar cancer-specific H3K4me3-BDs associated with hijacking of super-enhancers of other common oncogenes in B-cell (*MAF*, *MYC* and *FGFR3/NSD2*) and T-cell malignancies (*LMO2, TLX3* and *TAL1*). Our analysis suggests that H3K4me3-BDs can be created by super-enhancers and supports the new concept of *epigenomic translocation*, where the relocation of H3K4me3-BDs from cell identity genes to oncogenes accompanies the translocation of super-enhancers.

## Introduction

In healthy cells, many proto-oncogenes and tumour-suppressor genes coordinate to control cell proliferation. However, these essential genes can be activated or inactivated by different genomic alterations to their coding sequence, including nucleotide substitutions, gene amplification or loss, and gene fusion. Proto-oncogenes can also be converted to oncogenes without alterations to the protein-coding sequence; structural alterations can result in juxtaposition of proto-oncogenes and super-enhancers promoting their overexpression, a situation that is referred to as *enhancer adoption* (Lettice et al. 2011) or, more often, *enhancer hijacking* (Northcott et al. 2014; Beroukhim, Zhang, and Meyerson 2016; Weischenfeldt et al. 2017). Other structural alterations that disrupt local chromatin architecture can produce epigenomic activation of proto-oncogenes to resemble adjacent chromosomal neighbourhoods (Hnisz et al. 2016). However, little is known about the epigenomic consequences of enhancer hijacking on the deregulated oncogenes. To begin to understand how structural alterations activate proto-oncogenes, we have analysed the changes in chromatin states associated with super-enhancer translocation events in cancer.

Tri-methylation of lysine 4 on histone H3 (H3K4me3) is a chromatin modification classically associated with the promoters of transcriptionally active genes (Howe et al. 2017) and also present at some active enhancers (Pekowska et al. 2011; Shen et al. 2016; Hu et al. 2017; Russ et al. 2017; Q.-L. Li et al. 2019). Although H3K4me3 marks tend to show sharp 1– 2 kb peaks around promoters, some genes have broader regions of H3K4me3, also known as H3K4me3 broad domains (H3K4me3-BDs) that expand over part or all of the coding sequence of the gene (up to 20 kb) (Pekowska et al. 2010, 2011; K. Chen et al. 2015; Benayoun et al. 2014). These H3K4me3-BDs are associated with cell identity genes (Pekowska et al. 2010; Benayoun et al. 2014) and cell-specific tumour-suppressor genes (K. Chen et al. 2015), where they favour transcriptional consistency and increased expression (Benayoun et al. 2014). It has been observed that some tumour-suppressor genes show a narrower breadth of these domains in cancer, associated with their downregulation in malignant cells (K. Chen et al. 2015).

Super-enhancers were originally defined as enhancers with unusually high levels of certain transcriptional co-activators (Whyte et al. 2013; Lovén et al. 2013) and a median size larger than 8 kb (Pott and Lieb 2015). Recent studies of 3D genome networks suggest that the presence of H3K4me3-BDs in genes is associated with increased interactions with active super-enhancers (Cao et al. 2017; Thibodeau et al. 2017). More recently, Dhar and co-workers have shown that the histone methyl-transferase, MLL4, is essential for the methylation of both super-enhancers and H3K4me3-BDs, and proposed a hypothetical model whereby MLL4 would be essential to maintain the interaction of super-enhancers and tumour-suppressor genes with BDs (Dhar et al. 2018). However, the data from these previous studies could not address whether H3K4me3-BDs are formed as a consequence of super-enhancer activity.

The immunoglobulin (Ig) loci contain powerful super-enhancers to drive antibody formation and expression. The immunoglobulin heavy locus (*IGH*) has four regions encompassing: constant (C_H_), joining (J_H_), diversity (D_H_) and variable (V_H_) gene segments. The human C_H_ region encodes nine different Ig isotypes: IGHA2, IGHE, IGHG4, IGHG2, IGHA1, IGHG1, IGHG3, IGHD and IGHM (**Supplemental Fig. S1**). Complex regulatory and genomic rearrangements in the *IGH* locus are required to ensure that immunoglobulin transcripts containing only one of each of these gene segments are expressed in each B cell at the correct stage of B-cell differentiation. These natural processes predispose human Ig loci to inappropriate translocation events. As a result, these super-enhancer rich regions are commonly hijacked in B-cell malignancies providing the optimal model in which to address whether H3K4me3-BDs are formed as a consequence of super-enhancer activity.

## Results

### Identification of H3K4me3-BDs using the chromatin state model

H3K4me3-BDs have been defined in different ways, using size and height-ranked (Benayoun et al, 2014 and Chen et al. 2015, respectively) H3K4me3 peaks and, more recently, using a selection method based on two (intermediate and high) inflection points of ranked H3K4me3 peaks (Belhocine et al. 2021). However, all identified the enrichment of similar biological processes related to cell identity. In this study we defined H3K4me3-BDs and super-enhancers using the chromatin state model developed by Carrillo-de-Santa-Pau et al. 2017 using BLUEPRINT data of H3K4me3, H3K4me1, H3K27ac, H3K27me3, H3K9me3 and H3K36me3 (**Supplemental Tables S1–S3**). We defined H3K4me3-BDs as domains with H3K4me3 larger than 2 kb and compared them across different haematopoietic cell types (**Supplemental Results, Supplemental Table S4, Supplemental Fig. S2**). We observed shared H3K4me3-BDs in all of the cell types that were associated with basic cellular functions (**Supplemental Fig. S3D**). Only a small proportion of detected H3K4me3-BDs were cell-type exclusive (**Supplemental Fig. S2**), identifying cell identity genes (**Supplemental Fig. S3A–C**) as reported by the other studies mentioned above. With recent studies observing increased interactions between H3K4me3-BD and super-enhancers (Cao et al. 2017; Thibodeau et al. 2017), we also assessed the proximity of super-enhancers (defined as domains with H3K4me1 and H3K27ac larger than 5 kb) to genes marked with narrow promoter restricted H3K4me3 peaks or H3K4me3-BDs. In agreement with previous reports, we identified a significantly higher proportion of H3K4me3-BD having a proximal super-enhancer(s) within 100 kb in comparison to genes marked with narrow promoter restricted H3K4me3 peaks (**Supplemental Fig. S4**). The cell-type exclusive H3K4me3-BDs (**Supplemental Fig. S4D–F**) had proximal super-enhancer(s) more frequently than cell-type non-exclusive H3K4me3-BDs (**Supplemental Fig. S4A–C**).

### Cancer associated translocation events provide opportunities to investigate super-enhancer and H3K4me3-BD relationships

A common translocation in B-cell malignancies involves the proto-oncogene cyclin D1 (*CCND1*) at 11q13 and *IGH* at 14q32 (*IGH-CCND1*). To begin to understand how this translocation event leads to *CCND1* activation, we characterised the epigenomic landscape in these two regions from healthy and malignant B cells. To precisely define the location and activity of the human *IGH* super-enhancers and promoters and the chromatin dynamics of the *CCND1* locus in B cells, we used 108 chromatin state maps for different haematopoietic cell types built with over 700 ChIP-seq datasets from BLUEPRINT (Carrillo-de-Santa-Pau et al. 2017; Stunnenberg, International Human Epigenome Consortium, and Hirst 2016) (**Supplemental Tables S1–S3**). These data included samples from haematopoietic stem cells, four stages of healthy B-cell differentiation, plasma cells and both primary tumour cells and cell lines derived from four different B-cell haematological malignancies. We used seven healthy non-B-cell cell types (T cells, neutrophils, eosinophils, monocytes, macrophages, erythroblasts and megakaryocytes) as negative controls in which chromatin activity signals in the *IGH* locus are not predicted.

### Epigenomic cartography of the human IGH locus

Three super-enhancers have been previously described in the non-variable region of *IGH*: Eα2, Eα1 and Eμ (Mills et al. 1983; C. Chen and Birshtein 1997; Mills et al. 1997). However, the precise definition, location and activity of these regulatory regions throughout different stages of healthy human B-cell differentiation and human B-cell derived malignancies have not been previously investigated.

The three known B-cell specific active enhancers (high signal of both histone marks H3K4me1 and H3K27ac) (∼40 kb) were found within C_H_ region at the following genomic locations: Chr 14:106,025,200–106,056,800 (Eα2), Chr 14:106,144,200–106,179,400 (Eα1) and Chr 14:106,281,800–106,326,200 (Eμ). The Eμ enhancer showed dynamic changes associated with the B-cell development stage. A larger region appears active in naïve B cells before entering the germinal centre (GC), whereas in post-GC B cells the region of activity is reduced (class-switched memory B cells and plasma cells, ***Fig. 1***). A similar area of reduced activity was also observed in malignant samples derived from cells post-class-switch recombination, including diffuse large B-cell lymphoma (DLBCL), Burkitt lymphoma (BL) and multiple myeloma (MM) in comparison to pre-GC-like malignancies such as mantle cell lymphoma (MCL). We propose that a second, independent enhancer should be distinguished from Eµ based on the observation of a ∼10 kb gap between these two enhancer regions that is supported by high quality mapping. We refer to this as Eδ (Eδ at Chr 14:106,281,800–106,289,800 and Eμ at Chr 14:106,299,800–106,326,200) (***Fig. 1***). None of these four enhancer regions were detected in healthy myeloid and T cells.

**Figure 1:**
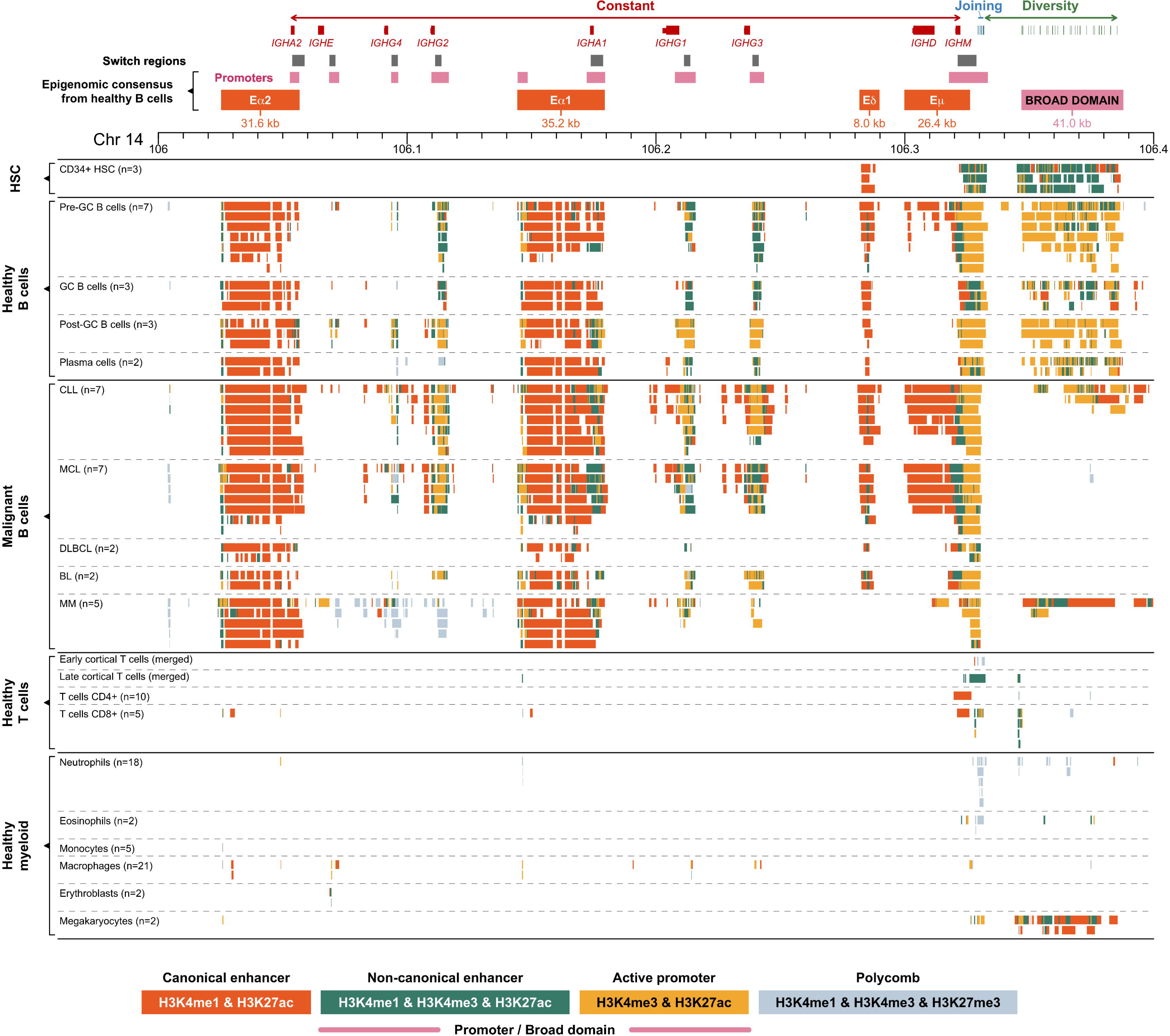
Genomic and epigenomic architecture of the *IGH* locus (14q32) in healthy and malignant human haematopoietic cells. Each panel represents collapsed cell-type specific signal of ChIP-seq chromatin states included in this study (**Supplemental Table S1**). AID motif clusters were detected as high enrichment of AID motifs (>200 of RGYW motifs per 2.5 kb). Abbreviations: HSC – Haematopoietic stem cells, GC – Germinal centre, CLL – Chronic lymphocytic leukemia, MCL – Mantle cell lymphoma, DLBCL – Diffuse large B-cell lymphoma, BL – Burkitt lymphoma, MM – Multiple myeloma.

### Epigenomic cartography of the human *CCND1* gene

The proto-oncogene *CCND1* is one of the most common *IGH* translocation partner genes in haematological malignancies, a hallmark of MCL (Vose 2017) and frequently observed in MM (Walker et al. 2013). The chromatin states in the *CCND1* promoter and immediate upstream region (***Fig. 2***) suggest that the Polycomb repressive complex may be implicated in the regulation of this cell cycle gene in healthy human B cells. When the promoter is inactive in non-proliferating GC B cells and terminally differentiated plasma cells, the Polycomb state covers the whole promoter and upstream region (Polycomb long, ***Fig. 2***). The Polycomb state is maintained upstream of the promoter in healthy proliferating haematopoietic cell types including pre-GC and post-GC B cells, some T cells, macrophages and eosinophils (Polycomb short, ***Fig. 2***).

**Figure 2:**
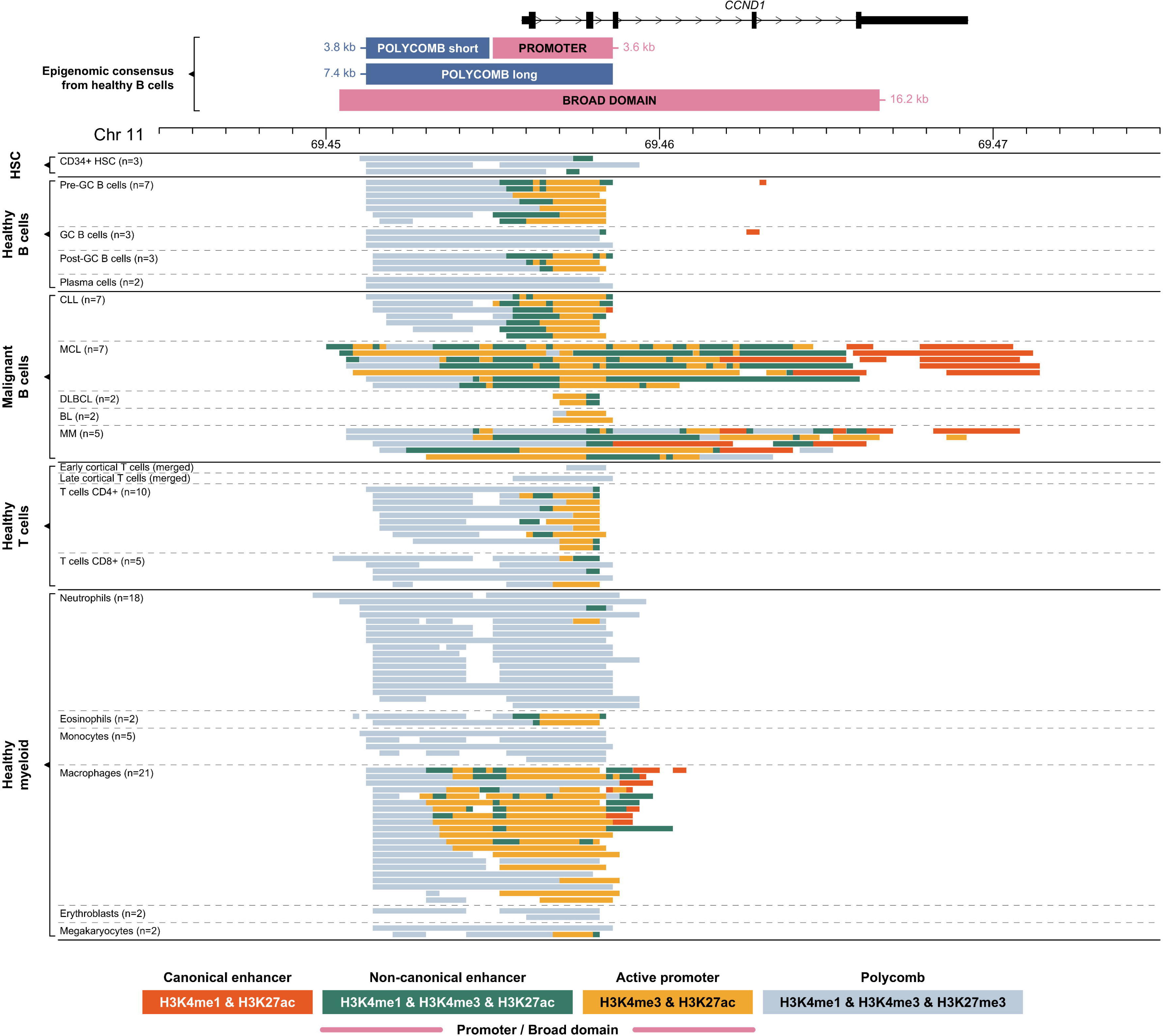
Genomic and epigenomic architecture of the *CCND1* locus (11p13) in healthy and malignant human haematopoietic cells. Each panel represents collapsed cell-type specific signal of ChIP-seq chromatin states included in this study (**Supplemental Table S1**). Abbreviations: HSC – Haematopoietic stem cells, GC – Germinal centre, CLL – Chronic lymphocytic leukemia, MCL – Mantle cell lymphoma, DLBCL – Diffuse large B-cell lymphoma, BL – Burkitt lymphoma, MM – Multiple myeloma.

We observed a different chromatin landscape for *CCND1* across four haematological malignancy subtypes (***Fig. 2***). In the majority of cases, the *CCND1* promoter shows an active state. Chronic lymphocytic leukaemia (CLL) samples show a similar pattern to healthy B cells, with the Polycomb state domain upstream of the active promoter. DLBCL and BL samples show a narrow H3K4me3 active promoter but lack the upstream Polycomb domain. All MCL and some MM samples have a larger active promoter/enhancer region extending upstream and downstream of the transcription start site, with most of the gene body containing H3K4me3 marks – a pattern that looks comparable to an H3K4me3-BD (Pekowska et al. 2010, 2011; Benayoun et al. 2014; K. Chen et al. 2015). Cytogenetic analysis for BLUEPRINT primary patient samples confirmed the presence of an *IGH-CCND1* rearrangement in all MCL patient samples. Although we were unable to confirm this for the MM samples, it is reported that 20% of MM patients will have an *IGH-CCND1* rearrangement (Avet-Loiseau et al. 2007). We therefore hypothesised that the juxtaposition of an *IGH* super-enhancer close to the coding region of *CCND1* results in the presence of an H3K4me3-BD over the coding region of the gene.

### Re-location of an H3K4me3-BD from IGH to CCND1 as a consequence of a hijacked super-enhancer

We discovered a B-cell specific H3K4me3-BD at genomic location, Chr 14:106,346,800-106,387,800 (41 kb), overlapping the D_H_ region of the *IGH* locus (***Fig. 1***). This element was characterised by high signal of both H3K4me3 and H3K27ac (typical for promoters), combined with occasional H3K4me1 high signal. This H3K4me3-BD is absent or significantly reduced in malignant B cells. These data suggest that the disappearance of the H3K4me3-BD from the *IGH* locus may be the consequence of *IGH* super-enhancer hijacking via genomic translocation events that are known to be present in the malignancy subtypes represented (**Supplemental Table S2 and S3**).

To further investigate the presence or absence of H3K4me3-BD as a result of *IGH* translocations, we studied the MM cell line U266 in which the *IGH* super-enhancer, Eα1, is inserted next to the *CCND1* proto-oncogene (Gabrea et al. 1999). This cell line provides the opportunity to analyse the consequence of the relocation of an isolated *IGH* super-enhancer. Using paired-end read targeted DNA sequencing, we precisely mapped the chromosomal changes at the *IGH* locus (**Supplemental Fig. S5**). We confirmed two chromosomal breakpoints that occur in *IGHE* and *IGHA* switch regions and result in the cut and paste of the *IGH* Eα1 super-enhancer into Chromosome 11, approximately 12 kb upstream of *CCND1* (***Fig. 3***). The vast majority of the H3K4me3-BD observed over the D_H_ region in healthy B cells is absent from this translocated cell line. A small region of H3K4me3 is still observed over the D_H_ region, perhaps due to the presence of *IGH* Eα2 super-enhancer that enables the expression of the IgE monoclonal immunoglobulin produced by U266 cells (Hellman et al. 1988). Importantly, a cancer-specific H3K4me3-BD is observed covering most of the *CCND1* gene body (***Fig. 3***). This observation suggests the presence of an *epigenomic translocation* of the H3K4me3-BD from *IGH* to the *CCND1* locus, present exclusively in MCL and MM samples (***Figs. 1–2***). This epigenomic translocation can result from both errors in VDJ recombination (Stamatopoulos et al. 1999) as observed in MCL, and class-switch recombination (Walker et al. 2013) as observed in MM. An aberrant H3K4me3-BD over the adjacent gene *MYEOV* was also observed in the U266 cell line (**Supplemental Fig. S6**). This suggests that the inserted Eα1 super-enhancer has a bi-directional effect, generating an H3K4me3-BD over another adjacent gene.

**Figure 3:**
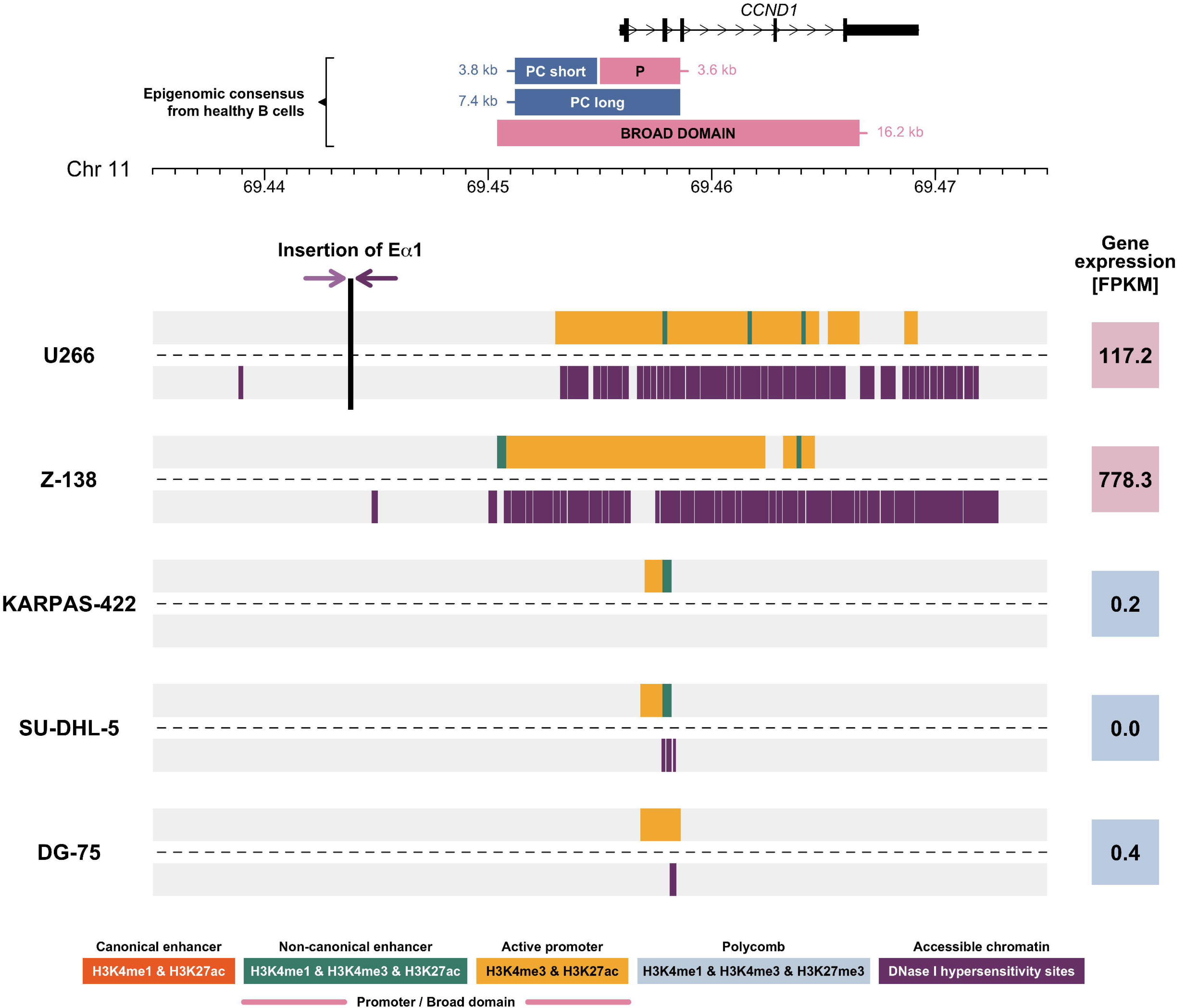
Chromatin landscape of the *CCND1* locus in five cell lines derived from B-cell haematological malignancies. Upper line of each cell line (**Supplemental Table S3**) represents the selected ChIP-seq chromatin states (**Supplemental Table S1**) and the lower line shows DNase I hypersensitivity sites for the non-variable region of the *IGH* locus. Vertical black line in U266 mark position of the inserted Eα1 super-enhancer. The breakpoints within the *IGH* locus are characterised in **Supplemental Fig. S5**. Shades of purple arrows symbolise translocation orientation. Numbers in coloured squares (red denotes high expression and blue low expression) show *CCND1* expression detected using RNA-seq in Fragments Per Kilobase of transcript per Million mapped reads (FPKM). Abbreviations: PC – Polycomb, P – Promoter.

### Relocation of H3K4me3-BD is associated with increased chromatin accessibility and transcription

To understand the effect of the epigenomic translocation on chromatin accessibility we used BLUEPRINT DNase I hypersensitivity data available for five cell lines derived from B-cell malignancies (**Supplemental Table S3**). Two of the cell lines had an *IGH-CCND1* rearrangement including U266 (already described above) and MCL cell line, Z-138. We observed the H3K4me3-BD and strong DNase I hypersensitivity signals encompassing the entire coding region of the *CCND1* gene in both cell lines (***Fig. 3***). The increased chromatin accessibility extended beyond the coding region of *CCND1*. This pattern is associated with an increase in *CCND1* expression in U266 and Z-138 (***Fig. 3***). The presence of the H3K4me3-BD, the strong DNase I hypersensitivity signal and increased transcript levels for *CCND1* were not observed in the remaining three cell lines without the *IGH-CCND1* rearrangement (***Fig. 3***). Whilst *MYEOV* showed an H3K4me3-BD, increased chromatin accessibility and expression in U266, this was not observed in Z-138 whereby the reciprocal translocation results in *MYEOV* being retained on the derived Chromosome 11, with the relocation of *CCND1* to Chromosome 14 juxtaposing it to all three *IGH* super-enhancers (Guikema et al. 2005; Rico et al., 2021).

### Cancer-specific H3K4me3-BDs are associated with immunoglobulin translocations in multiple myeloma

To investigate whether the epigenomic translocation of the *IGH* H3K4me3-BD can occur over additional proto-oncogenes in B-cell malignancies, we generated ChIP-seq data for the same six histone marks for an additional two MM cell lines (KMS11 and MM1S) and seven MM patients whose diagnostic samples were engrafted in murine models (patient-derived xenografts, or PDXs). Each cell line had a complex rearrangement identified by targeted sequencing of the *IGH* locus involving multiple proto-oncogenes (**Supplemental Table S3**). 10x genome sequencing detected in patients P1 and P2 the second most common *IGH* translocation in MM, t(14;16)(q32;q23), involving the transcription factor *MAF*. Patients P3 and P4 had *IGH* translocations t(4;14)(p16;q32), involving *FGFR3* and *NSD2* with the remaining three patients (P5–7) having no detectable translocation involving any of the Ig loci (**Supplemental Table S5**). No cell lines or patients had an *IGH-CCND1* rearrangement detected by targeted or genome sequencing therefore, as expected, we did not observe an aberrant H3K4me3-BD or an increase in mRNA transcript levels for *CCND1* in these samples (***Fig. 4A***). This suggests that epigenomic translocations of H3K4me3-BDs over *CCND1* are a direct result of a specific genomic translocation event.

**Figure 4:**
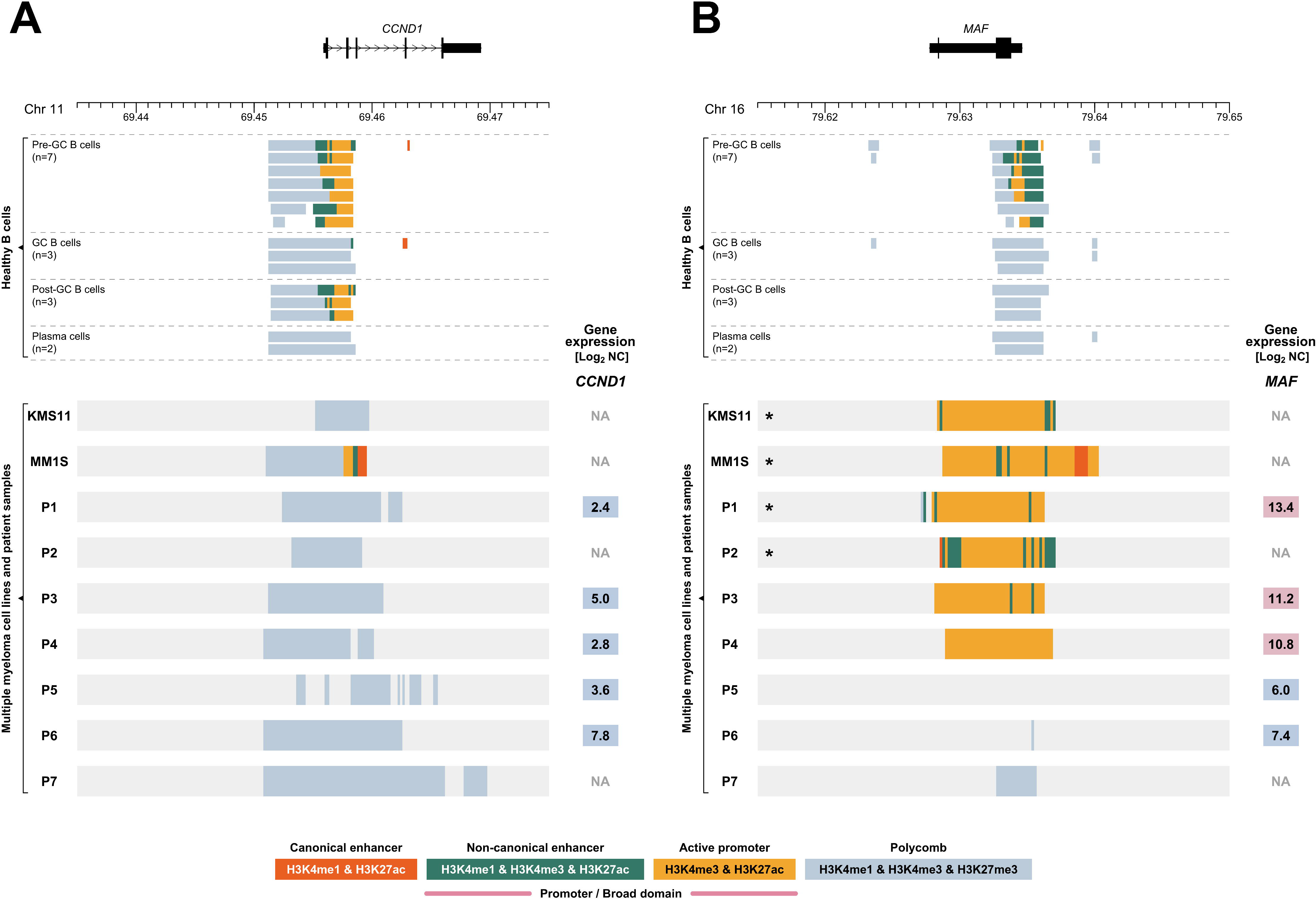
Chromatin landscape of the *CCND1* (11q13) and *MAF* (16q23) loci in healthy human B cells and multiple myeloma samples. Upper panels show selected ChIP-seq chromatin states (**Supplemental Table S1**) of *CCND1 (A)* and *MAF (B)* loci in BLUEPRINT healthy B cells. Each line of lower panel represents the ChIP-seq chromatin states for myeloma cell lines KMS11 and MM1S and seven multiple myeloma patients (P1-P7, patient derived xenograft material) for *CCND1 (A)* and *MAF (B)* loci. Numbers in coloured squares (red denotes high expression and blue low expression) show gene expression detected by RNA-seq and displayed as Log_2_ normalised counts (Log_2_ NC). GC – Germinal centre, *sample contains chromosomal translocation involving the displayed region, described in detail in **Supplemental Tables S3 and S5**.

In the two cell lines and four of the patient samples (P1–4) we observed an H3K4me3-BD over the coding region of *MAF* that was coupled with higher *MAF* expression observed by RNA-seq (***Fig. 4B***). This was associated with the presence of the *IGH-MAF* rearrangement in the two cell lines and patients P1 and P2. Our targeted sequencing identified the involvement of *MYC* as well in the complex translocations observed in KMS11 and MM1S cell lines (**Supplemental Table S3**). We observed a Polycomb chromatin state over *MYC*, but the analysis of each histone modification separately identified an H3K4me3-BD over *MYC* in the two cell lines (**Supplemental Figs. S7A** and **S8A**), possibly reflecting repression of one allele of *MYC* and activation of the other due to being juxtaposed to an IGH super-enhancer (Affer et al. 2014).

In the remaining two patients (P3, P4), we were unable to detect a genomic rearrangement of a super-enhancer in close proximity *to MAF* however, we see overexpression of *MAF* in association with an H3K4me3-BD (***Fig. 4B***). These two patients have the translocation t(4;14)(p16;q32) that involves *FGFR3* and *NSD2* genes. We observed an H3K4me3-BD and increased transcript levels for *FGFR3* and *NSD2* and a Polycomb chromatin state without H3K4me3-BD was observed over *MYC* coupled with low *MYC* expression in both patients (**Supplemental Figs. S7B** and **S8B**).

In summary, we have observed the appearance of H3K4me3-BDs over a variety of proto-oncogenes when they are involved in the hijacking of an *IGH* super-enhancer. Whilst the chromatin state model (Carrillo-de-Santa-Pau et al. 2017) identified BDs over *CCND1* and *MAF*, it preferentially selected a Polycomb chromatin state for both *MYC* and *FGFR3* in some samples. The presence of both H3K27me3 and H3K4me3 domains over these genes potentially highlights the repression of the second allele (Nag et al. 2015), a mechanism the cell can use to control the level of overexpression. With the H3K4me3-BD signature observed over *MYC* in both healthy and malignant B cells in this study and that of Belhocine and co-workers (Belhocine et al. 2021), future work will be required to unravel the epigenetic impact upon this locus after different super-enhancer translocations.

### Super-enhancer-driven H3K4me3-BDs as a wider phenomenon associated with oncogene deregulation in haematological malignancies

T-cell receptor loci (*TRA, TRD, TRB, TRG*) are also regulated by super-enhancers, undergo similar inherent somatic rearrangement events and are involved in translocation events in T-cell acute lymphoblastic leukaemia (T-ALL) (**Supplemental Fig. S9**) (Larmonie et al. 2013). To investigate whether these loci also have the ability to generate H3K4me3-BDs over proto-oncogenes involved in translocation events, we studied the KOPT-K1 T-ALL cell line that results in the juxtaposition of *LMO2* and the super-enhancer of the *TRA/TRD* locus as a result of the translocation, t(11;14)(q13;q11). The Polycomb chromatin state was observed in healthy early and late cortical T cells and *CD4* and *CD8A* expressing T cells (***Fig. 5A***). We indeed observed an aberrant H3K4me3-BD over the coding region of *LMO2* in KOPT-K1 suggesting that T-cell receptor super-enhancers can also generate H3K4me3-BD that results in proto-oncogene activation and overexpression (***Fig. 5A***) (Sanda et al. 2012). An H3K4me3-BD was not observed over *LMO2* in two additional cell lines that do not have rearrangements involving this locus.

**Figure 5.**
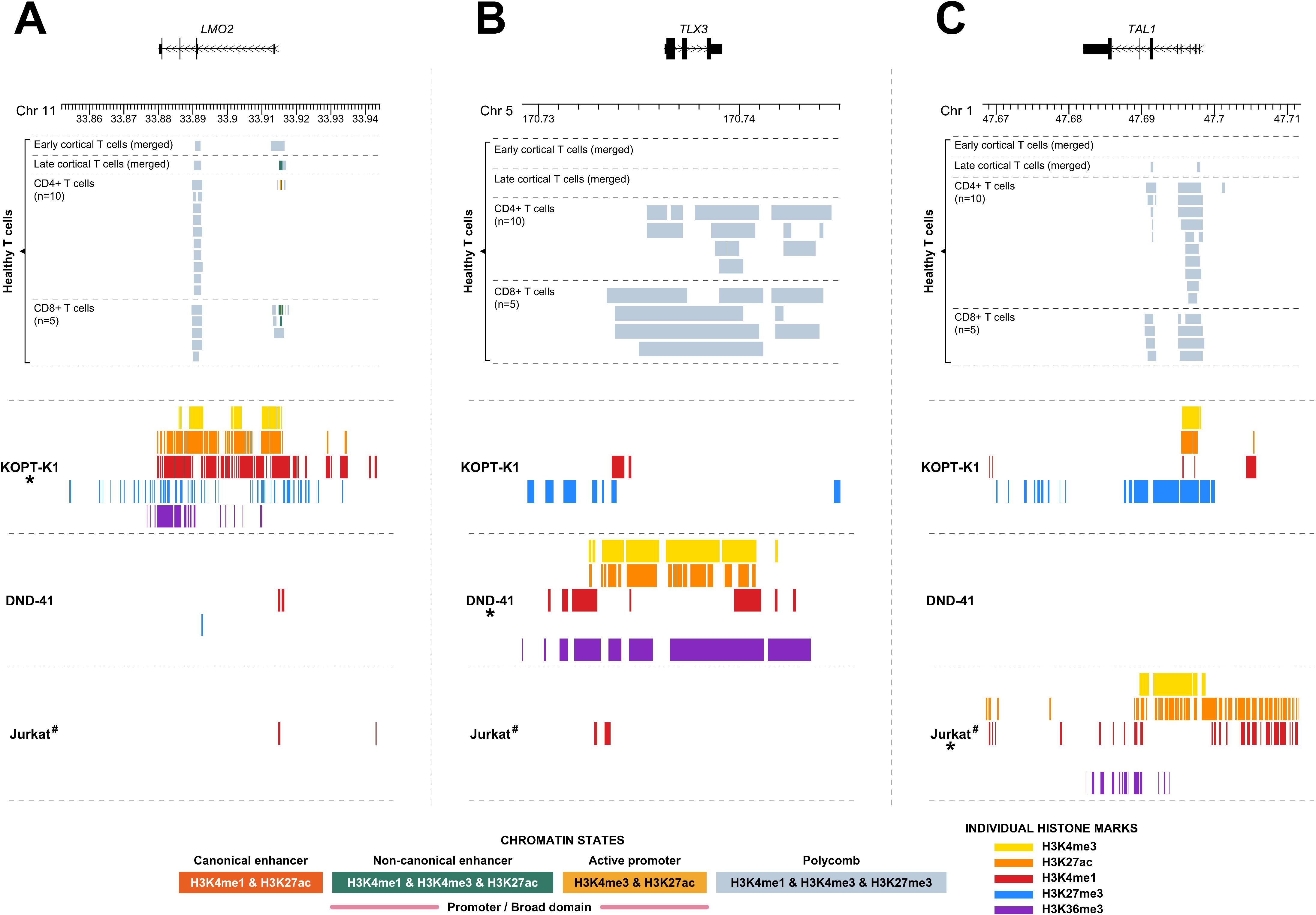
Chromatin landscape of the *LMO2* (11p13), *TLX3* (5q35) and *TAL1* (1p33) loci in healthy mature T cells and three cell lines derived from T-cell haematological malignancies. Upper panels show selected ChIP-seq chromatin states (**Supplemental Table S1**) of *LMO2 (A)*, *TLX3 (B)* and *TAL1 (C)* loci in BLUEPRINT healthy mature T cells. Each line of lower panels represents peaks of individual histone marks, separately for each cell line. *cell line contains chromosomal aberration involving the displayed region as follows: KOPT-K1 – *LMO2*-*TRA*/*TRD*, DND-41 – *TLX3*-*BCL11B*, and Jurkat – 12 bp insertion upstream of the *TAL1* gene (for details see **Supplemental Table S3**). missing data for histone mark H3K27me3 in Jurkat cell line.

To investigate the wider involvement of H3K4m3-BDs in T-ALL we considered the T-cell identity (Ha et al. 2017) and tumour suppressor gene, *BCL11B* that is rearranged in T-ALL (Liu et al. 2017). Since this gene is known to be regulated by a downstream super-enhancer element (**Supplemental Figs. S10** and **S11A**) (Nagel et al. 2007; L. Li et al. 2013), we questioned whether the relocation of this region from Chromosome 14 band q32 next to *TLX3* on Chromosome 5 band q35 would result in the appearance of an aberrant H3K4me3-BD over the coding region of the gene. Using publicly available ChIP-seq data for T-ALL cell line DND-41 (Knoechel et al. 2014) we observed high signal for H3K27ac, H3K4me3 and H3K4me1 encompassing the entire coding region of *TLX3* and non-coding regions either side (***Fig. 5B***). This correlated with a high transcript level when compared to healthy cells (Petryszak et al. 2016). We did not observe the presence of these histone marks in normal healthy T cells and cell lines without rearrangements of these loci (***Fig. 5B***).

Next we wanted to assess whether *de novo* super-enhancers, generated by somatic nucleotide insertions, could also generate aberrant H3K4me3-BD. An aberrant H3K4me3-BD was observed across the *TAL1* gene in the T-ALL cell line Jurkat (***Fig. 5C***) where a 12 bp insertion generates a super-enhancer element upstream in the gene (Mansour et al. 2014; Navarro et al. 2015). No super-enhancer and/or H3K4me-BD activity was observed upstream of the *TAL1* gene in healthy T cells (**Supplemental Fig. S12**). To provide mechanistic data to support the generation of H3K4me3-BD by super-enhancer relocation, we used our previously CRISPR-Cas9 engineered T-ALL cell line, PEER, that contains the same 12 bp insertion upstream of the *TAL1* gene resulting in *TAL1* overexpression (Navarro et al. 2015). H3K4me3 ChIP-seq confirmed broader H3K4 methylation upstream and over the gene body of *TAL1* in the engineered cells compared to wildtype (**Supplemental Fig. S13**).

In summary, we found that the appearance of H3K4me3-BDs is associated with cancer-specific super-enhancer activation in both B-cell and T-cell derived haematological malignancies. Our data suggest that H3K4me3-BDs could be a signature of super-enhancer targets in general and a useful marker to identify deregulated genes affected by the hijacking of super-enhancers in cancer.

## Discussion

We aimed to understand how the juxtaposition of super-enhancers and proto-oncogenes results in exceptionally high, but restricted levels of oncogene expression. For this, we first dissected the epigenomic landscape of two loci that are commonly translocated in haematological malignancies, *CCND1* at 11q13 and the *IGH* locus at 14q32. We have shown that genomic relocation of the *IGH* E 1 super-enhancer alters the location of an H3K4me3-BD in rearranged malignant B cells. The presence of broad H3K27ac domains over *CCND1* have previously been associated with t(11;14) translocations in MM cell lines including U266 (Jin et al. 2018) but this study did not use H3K4me3 data. These domains were described as *de novo* (genic) super-enhancers associated with *CCND1* overexpression but they are likely showing the H3K4me3-BDs we describe here (defined as chromatin states with high levels of H3K4me3 and H3K27ac. We also observed H3K4me3-BDs over additional proto-oncogenes rearranged in B-cell and T-cell malignancies including the involvement of gene specific and *de novo* super-enhancers. H3K4me3-BDs are associated with high levels of stable transcription of cell identity and tumour suppressor genes (Pekowska et al. 2010, 2011; Benayoun et al. 2014; K. Chen et al. 2015) and we show here, for the first time, that they can be a feature of oncogene activation by translocated super-enhancers (as a result of a genomic rearrangement).

We analysed a total of fourteen haemato-oncology samples, newly generated data for seven PDX samples and seven patient-derived cell lines. Twelve of these cases were positive for a genomic abnormality involving hijacking of super-enhancers and proto-oncogene activation (remaining two did not have these rearrangements). All twelve samples showed an H3K4me3-BD over the oncogene specific to a genomic rearrangement, confirming our hypothesis about the association of H3K4me3-BD and super-enhancers. In three of these twelve samples, additional H3K4me3-BDs were observed over other oncogenes not involved in a genomic rearrangement, however, there was no evidence of super-enhancer translocations at these loci. This could be due to unidentified chromosomal abnormalities involving other super-enhancers or by the existence of additional mechanisms associated with H3K4me3-BD formation. This is in agreement with our complementary publication by Belhocine and co-workers (Belhocine et al. 2021) where we observed the generation of H3K4me3-BD over many cancer-related genes suggesting the presence of additional mechanisms able to lay down H3K4me3-BDs over coding and non-coding regions of the genome. With super-enhancers hijacking essential transcription factors (*MYC*) and chromatin remodelers (*NSD2*) it may not be surprising that the increased expression and presence of the translated protein could indirectly drive the appearance of H3K4me3-BD over other cancer associated genes.

Based on these data, we present a model of *epigenomic translocation*, a consequence of hijacking of super-enhancers. In this model a super-enhancer driven wild-type H3K4me3-BD “re-locates” to a target proto-oncogene as the result of a genomic rearrangement leading to oncogene activation (***Fig. 6***).

**Figure 6:**
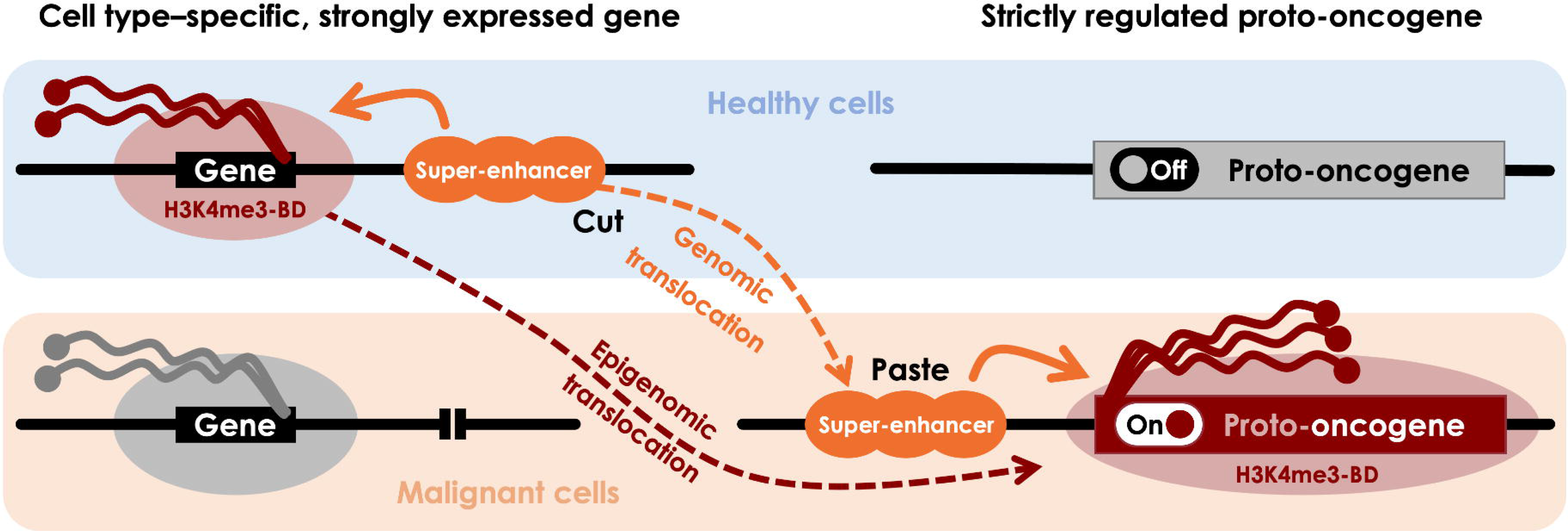
Scheme of epigenomic translocation model. In healthy cells, a cell identity or tumour suppressor gene is expressed by regulation of a super-enhancer that generates an H3K4me3 broad domain (H3K4me3-BD), allowing rapid and consistent activation of the transcriptional machinery (upper left). Proto-oncogenes are strictly regulated to control important physiological processes including the cell cycle (upper right). During a genomic translocation event, a super-enhancer is juxtaposed close to a proto-oncogene resulting in oncogenic activation (bottom right). Re-location of the super-enhancer brings the transcriptional machinery including the H3K4me3-BD. This epigenomic signature disappears at the original locus (bottom left).

Our model implies that H3K4me3-BDs can be generated by super-enhancers in their wildtype neighbourhood. The H3K4me3 writer MLL4 is essential for the maintenance of both H3K4me3-BDs and super-enhancers, but not for narrow-peak promoters and standard enhancers (Dhar et al. 2018). We have recently shown that inhibition of transcriptional elongation machinery by targeting CDK7 and DOT1L, or by inhibiting H3K4me3 demethylation with the use of PBIT inhibitors, can preferentially impact H3K4me4-BD and cellular viability (Belhocine et al. 2021). Initially, a strong emphasis was placed on the distinction between H3K4me3-BDs and super-enhancers (Benayoun et al. 2014) even if they were later shown to be close or sometimes overlapping in the linear genome (Cao et al. 2017; Chen et al. 2015) and in 3D (Thibodeau et al. 2017). Whilst super-enhancers may be responsible for generating broad domains, very few studies have addressed the length and maintenance of these regions.

Recent simulations and experimental data suggest that very active enhancers may induce high levels of gene expression by global chromatin decompaction that may actually increase the distance between promoter and enhancer regions (Buckle et al. 2018). We have observed that the presence of the H3K4me3-BD over *CCND1* coincides with high chromatin accessibility of the whole gene body. If super-enhancers induce unusual chromatin decompaction and access to DNA, this could explain how the target genes of oncogenic translocations can show such high levels of gene expression. The effects of the super-enhancer could be restricted within the boundaries of the topologically associated domains (TADs) where the hijacked oncogene and super-enhancer reside (Gong et al. 2018). Indeed, it has been suggested that one reason why TAD boundaries are conserved across species (Lazar et al. 2018; Krefting, Andrade-Navarro, and Ibn-Salem 2018) could be that they prevent undesirable enhancer hijacking (Fudenberg and Pollard 2019). In related work, we have used the highly-predictive heteromorphic polymer (HiP-HoP) model (Buckle et al. 2018) to predict chromatin conformations at the proto-oncogene *CCND1* in healthy and malignant B cells (Rico et al, 2021). Our simulations of U266 and Z-138 cancer cell lines harbouring *IGH-CCND1* rearrangements predict extensive changes in enhancer-promoter interactions, providing additional evidence that the downstream chromatin remodeling is essential for oncogene overexpression. In U266 cells the TAD structure was unchanged with the *IGH* super-enhancer providing strong predicted interactions with the promoter and gene body of *CCND1* (Rico et al, 2021). In Z-138 cells, the reciprocal translocation generated a new oncogenic TAD on Chromosome 14 with the strongest interactions observed between the *IGH* Eμ and Eδ super-enhancers, again with the promoter and gene body of *CCND1* (Rico et al. 2021). However, it is important to note that the H3K4me3-BD is not necessarily causing the oncogene overexpression, and it could actually be a consequence of the super-enhancer driven overexpression of the oncogene (Howe et al. 2017).

The epigenomic translocation scenario is a simple yet powerful conceptual model that extends the original enhancer adoption/hijacking model (Northcott et al. 2014; Beroukhim, Zhang, and Meyerson 2016; Lettice et al. 2011; Weischenfeldt et al. 2017) and allows inference of candidate genes deregulated as result of translocation events. Although we propose that some broad H3K4me3 domains are the consequence of active super-enhancers, future studies will be needed to elucidate the possible cooperation between genetic features, other epigenetic marks, the broad domain and the super-enhancer chromatin structures and how they maintain each other.

This work will hopefully pave the way for new therapeutic approaches based on chromatin remodelling to revert the local epigenome of super-enhancer activated proto-oncogenes, returning their expression back to wild-type levels. We anticipate the model proposed here will focus attention on the regulatory effects of different genomic rearrangements including translocations, identifying key oncogenes in each patient, and open exciting new avenues for novel diagnostic and therapeutic approaches.

## Methods

### ChIP-seq, DNase I hypersensitivity and RNA-seq of BLUEPRINT dataset

Previously processed ChIP-seq chromatin state BLUEPRINT samples (n=108) were used (**Supplemental Tables S2–S3**). This dataset comprised 85 samples from healthy donors (3 haematopoietic stem cells, 15 B-cell lineage, 17 T-cell lineage and 50 myeloid lineage) and 23 samples from patients (16 primary and seven cell lines) with different B-cell haematological malignancies. Five of these cell lines had DNase I hypersensitivity and RNA-seq data available (**Supplemental Table S3**). BLUEPRINT data were downloaded from ftp://ftp.ebi.ac.uk/pub/databases/blueprint/paper_data_sets/chromatin_states_carrillo_bu <u>ild37</u> (ChIP-seq chromatin states) and ftp://ftp.ebi.ac.uk/pub/databases/blueprint/data/homo_sapiens/GRCh37 (DNase I hypersensitivity and RNA-seq). Early and late cortical T-cell ChIP-seq chromatin states are accessible by https://github.com/guillaumecharbonnier/mw-cieslak2019/tree/master/src/hub. Detailed laboratory methods and data processing of BLUEPRINT experiments are described elsewhere (Fernández et al. 2016; Stunnenberg, International Human Epigenome Consortium, and Hirst 2016; Carrillo-de-Santa-Pau et al. 2017; Cieslak et al. 2020). Briefly, sequencing reads were mapped using BWA (ChIP-seq and DNase-seq, v0.5.9) or GEM mapper (Marco-Sola et al. 2012) (RNA-seq) to GRCh37 human genome assembly. ChromHMM (Ernst and Kellis 2012) (v1.10) was used to determine chromatin states, based on a combination of six histone modifications as follows: H3K4me1, H3K4me3, H3K9me3, H3K27me3, H3K27ac and H3K36me3 (Carrillo-de-Santa-Pau et al. 2017). Selected chromatin states used in this study are described in **Supplemental Table S1**. Presence of *IGH-CCND1* rearrangement in MCL samples was detected by conventional cytogenetics and fluorescence *in situ* hybridization at the Laboratory of Oncohematological Cytogenetics of Hospital Clinic Barcelona (Spain).

### Epigenomic consensus determined by BLUEPRINT ChIP-seq data

Epigenomic annotation of *IGH* (super-enhancers, promoters and H3K4me3-BD) and *CCND1* (promoter and Polycomb) loci was built using 15 BLUEPRINT B-cell/plasma-cell samples derived from healthy donors (**Supplemental Table S2**). Epigenomic consensus of super-enhancers and H3K4me3-BDs at *TRA/TRD* and *BCL11B* loci were assembled using 17 BLUEPRINT T-cells samples derived from healthy donors. MCL (JVM-2 and Z-138) and MM (U266) cell lines with known *IGH-CCND1* rearrangements (**Supplemental Table S3**) were used to determine the relocated *CCND1* H3K4me3-BD.

Segmentation files of relevant samples were filtered for the subset of chromatin states (Carrillo-de-Santa-Pau et al. 2017) of interest as follows: active canonical enhancer (state 9) for super-enhancers, repressed Polycomb regulatory region (state 7) for Polycomb, and a combination of active promoter (state 11) and active non-canonical H3K4me3 enhancer (state 10) for promoters and H3K4me3-BDs. For each of these epigenomic elements, subsets were intersected and merged (both BEDTools v2.27.1) into continuous regions with skipping gaps smaller than 1 kb. Super-enhancers were called as regions bigger than 2.5 kb at *IGH* (overlapping the C_H_ and *IGH* intronic region), *TRA/TRD* (constant and joining regions) and *BCL11B* (from the gene position up to ∼1 Mb downstream) loci.

Promoters were determined as a segment bigger than 2.5 kb within the expected promoter positions of *IGH* (C_H_ and intronic region) and *CCND1* genes. H3K4me3-BDs were scanned at *IGH* C_H_, J_H_ and D_H_, and *CCND1*, *TRA/TRD* and *BCL11B* loci, with a minimum size of 15 kb. Polycomb at the *CCND1* locus was obtained as regions upstream and overlapping with the promoter.

### Genome-wide H3K4me3-BDs and super-enhancers co-occurrence

We used an epigenomic consensus approach to test whether H3K4me3-BDs co-occur with super enhancers at the genome-wide level in the BLUEPRINT dataset (**Supplemental Tables S2–S3**). For this analysis, the proximity of H3K4me3-BDs (≥2 kb) to super-enhancers were compared (≥5 kb) to a control group of small promoters (1 kb, overlapping the transcription start site). Super-enhancers overlapping directly H3K4me3-BD and/or genes with H3K4me3-BD were excluded from the analysis. Also, promoters in genes with H3K4me3-BD in alternative transcription start site were excluded. Genomic distances between H3K4me3-BD/promoters and super-enhancers up to 5 Mb were analysed by 100 kb bins. Gene enrichment for biological processes was performed using R package clusterProfiler v3.18.0. To determine epigenomic cell type specificity, H3K4me3-BDs were tested for overlaps and coverage of an active chromatin state background for each H3K4me3-BD was calculated. A threshold <10% was used for low active chromatin state background coverage.

### Targeted DNA sequencing of myeloma cell lines

DNA of myeloma cell lines U266, KMS11 and MM1S (**Supplemental Table S3**) were sequenced using targeted high-throughput sequencing covering the following genomic regions: 1) 4.2 Mb; extended coverage for chromosomal abnormality detection within *IGH*, *IGK*, *IGL* and *MYC* (4.2 Mb); and 2) 0.6 Mb, exonic regions of 127 myeloma specific genes for mutation analysis and 27 additional regions for efficient data normalisation. GRCh37 human genome assembly was used for sequence mapping using BWA-MEM (v0.7.12). Chromosomal rearrangements were called using Manta v.0.29.6 (Mikulasova et al. 2019; X. Chen et al. 2016). Detailed methods are described previously (Mikulasova et al. 2019).

### Myeloma patient derived xenografts

Patient derived xenografts were generated by passaging primary patient CD138+ selected cells through the previously described SCID-rab myeloma mouse model (Yata and Yaccoby 2004; Mikulasova et al. 2019). Detailed methods are described previously (Mikulasova et al. 2019). Seven patient derived xenografts were used in this study (**Supplemental Table S5**).

### Whole genome sequencing of patient derived xenograft samples

DNA from seven patient derived xenograft samples were sequenced using phased whole genome sequencing (10x Genomics, Pleasanton, CA, USA) at Hudson Alpha (Huntsville, AL, USA). Long Ranger (10x Genomics) pipelines were used for data processing, including alignment to GRCh38 genome assembly and structural variant calling. Germline controls were used to distinguish somatic abnormalities and chromosomal breakpoints of detected translocations (**Supplemental Table S5**) were manually inspected.

### ChIP-seq of myeloma cell lines and patient derived xenograft samples

ChIP-seq was performed as previously reported (Mikulasova et al. 2019), for the myeloma cell lines KMS11 and MM1S as well as seven myeloma patient derived xenograft samples. ChIP-seq for the histone marks H3K4me1, H3K4me3, H3K9me3, H3K27me3, H3K27ac, and H3K36me3 (Active Motif, Carlsbad, CA, USA) were included in this study. Controls without antibody input were performed to ensure data quality. GRCh38 human assembly was used for alignment.

ChIP-seq chromatin states were determined by ChromHMM (v1.20), the model used in the BLUEPRINT dataset (Carrillo-de-Santa-Pau et al. 2017). Additionally, the peaks of individual histone marks were detected using MACS2 (v2.2.5) by a pipeline available at BLUEPRINT Data Coordination Center Portal (Fernández et al. 2016). For direct comparison of this data and BLUEPRINT data, The liftOver (University of California, Santa Cruz, CA, USA) tool was used to convert chromatin state segments and histone peaks within the genomic regions included in this study from GRCh38 to GRCh37 genome assembly. No segment was lost during this conversion.

### RNA-seq of patient derived xenograft samples

RNA-seq was performed using 100 ng total RNA with genomic DNA removal using the TURBO DNA-free kit (Ambion, Austin, TX, USA). RNA was prepared using the TruSeq stranded total RNA Ribo-Zero gold kit (Illumina, San Diego, CA, USA) and libraries were sequenced using 75 bp paired end reads on a NextSeq 500 (Illumina). RNA-seq data was analysed as previously reported (Mikulasova et al. 2019). Briefly, raw data were aligned to the human genome assembly GRCh38 with gene transcript quantification being processed by Star (v2.5.1b) and Salmon (v0.6.0) algorithms. Read counts per gene were read into R and using the DESeq2 (v1.20.0) R library, normalised across samples and the log_2_ expression calculated.

### ChIP-seq of T-cell acute lymphoblastic leukemia cell lines

Publicly available ChIP-seq data accessible by the numbers GSE54379 and GSE65687 at National Center for Biotechnology Information (NCBI, Bethesda, MD, USA) Gene Expression Omnibus (GEO) were used for histone modification analysis of three T-ALL cell lines KOPT-K1, DND-41 and Jurkat (**Supplemental Table S3**). Raw files were downloaded using the associated NCBI Sequence Read Archive (SRA) accession and converted to FASTQ files using NCBI fasterq-dump (SRA Toolkit v2.9.6-1). Appropriate sequencing runs were merged and sequences were mapped to GRCh37 genome assembly using BWA (v0.7.17), followed by processing to MACS2 (v2.2.5) peaks, using a pipeline available at BLUEPRINT Data Coordination Center Portal (http://dcc.blueprint-epigenome.eu/). Histone marks H3K4me1, H3K4me3, H3K27me3, H3K27ac, and H3K36me3 in all three cell lines were used in this study, with the exception of H3K27me3 in Jurkat as this was not available.

Additionally, wild type and CRISPR-Cas9 edited (upstream 12 bp insertion known from Jurkat cell line) PEER cell line were included. The genome editing method was published elsewhere (Navarro et al. 2015). ChIP-seq for H3K4me3 and H3K27ac in PEER cell line was generated using MicroPlex Library Preparation Kit (Diagenode, Liege, Belgium) according to the BLUEPRINT protocol (http://dcc.blueprint-epigenome.eu/#/md/methods). Prepared libraries were sequenced in paired-end 50+30bp mode using the NextSeq 500/550 (Illumina) according to manufacturer’s instructions. FASTQ files were processed by BLUEPRINT pipeline as T-ALL cell lines above. Raw read counts were collected using Genomic Analysis Toolkit v4.2 in 200bp bins and normalized by average number of reads per a bin.

### Statistical analysis and graphical output

Statistical analysis was performed using R v4.0.3 (R Core Team. 2020). P values <0.05 were considered statistically significant. Figures involving data and annotation alignment to the human genome were generated using karyoploteR (v1.16.0) R package.

### Human genome assembly statement

This study focuses on human genome loci where differences between hg19/GRCh37 and hg38/GRCh38 genome assemblies would not affect the results and conclusions.

### Data access

The DNA sequencing data for U266, KMS11 and MM1S myeloma cell lines generated in this study have been submitted to the NCBI BioProject database (https://www.ncbi.nlm.nih.gov/bioproject/) under accession number PRJNA635269. All raw and processed ChIP-seq data for KMS11 and MM1S myeloma cell lines as well as PEER T-ALL cell line have been submitted to the NCBI GEO (https://www.ncbi.nlm.nih.gov/geo/) under accession number GSE151556. All high-throughput sequencing data for multiple myeloma patient derived xenografts have been submitted to the European Genome-phenome Archive (https://ega-archive.org/) under accession number EGAS00001005684.

## Author contributions

L.J.R. and D.R. conceived and designed the study; A.E.C., S.H., B.A.W., D.R. and L.J.R supervised the research; A.M. led the data analysis and interpreted results with L.J.R and D.R.; A.M., D.K., M.T., N.K., K.T.M.F., C.A. and C.A.B., and processed and analysed genomic, epigenomic and chromatin accessibility data; C.A., S.Y., F.R., M.Z., S.T., C.S, G.J.M. and B.A.W. were responsible for collection and generation of myeloma data; A.C., V.A. and S.S. helped with T-cell data collection and interpretation, and generated ChIP-seq data of the PEER cell line; A.M., D.R. and L.J.R took the lead in writing the manuscript; all authors discussed the results, wrote, reviewed and approved the manuscript.

## Acknowledgements

This work was supported by the Wellcome Trust (206103/Z/17/Z to D.R. and 207556/Z/17/Z to S.H.), the Leukemia & Lymphoma Society (to G.J.M. and B.A.W.) and Kidscan Children’s Cancer Research and The Little Princess Trust (to L.J.R). S.H. would also like to thank the Newcastle upon Tyne Hospitals NHS Charities, the NIHR Newcastle Biomedical Research Centre and the Sir Jules Thorn Charitable Trust (12/JTA). S.S. and V.A. would also like to thank the Institut National du Cancer (PLBIO018-031 INCA_12619). L.J.R and D.R. both had Newcastle University Research Fellowships and A.M. a Newcastle University Faculty Fellowship.

