## Supplemental materials for "Epigenomic translocation of H3K4me3 broad domains over oncogenes following hijacking of super-enhancers"

[8]Equipe Labellisée Ligue Contre le Cancer

[9]SUPA, School of Physics and Astronomy, University of Edinburgh, Edinburgh, UK

[10]Lymphocyte Signalling and Development Programme, Babraham Institute, Cambridge, UK

[11]Great North Children's Hospital, Newcastle upon Tyne Hospitals NHS Foundation Trust, UK

[12]Melvin and Bren Simon Comprehensive Cancer Center, Division of Hematology Oncology, Indiana University, Indianapolis, IN, USA

[*]Joint last authors

**Supplemental Results**

**Genome-wide H3K4me3-BD and super-enhancer co-occurrence**

A total of 59,097 (15,443 non-overlapping) H3K4me3-BDs were detected at genome-wide level using BLUEPRINT chromatin state data divided into the following haematopoietic cell types: haematopoietic stem cells (HSC), B cells, T cells, myeloid cells, MCL, CLL and MM. H3K4me3-BDs were classified into categories based on epigenomic specificity using their overlaps and active chromatin background (**Supplemental Fig. S2**). A total of 31, 302, 144 and 1186 H3K4me3-BDs were exclusively detected in HSC, B cells, T cells and myeloid cells, respectively. This indicates that the vast majority of H3K4me3-BDs remain stable during haematopoiesis.

We confirm that cell-type exclusive H3K4me3-BDs are associated with cell-identity genes (**Supplemental Fig. S3**). B-cell exclusive H3K4me3-BDs identified genes important in B-cell biology (for example immunoglobulin genes *IGH*, *IGK* and *IGL*, or *PAX5*, *MS4A1* and *CD22*). Similarly, T-cell identity genes were enriched within T-cell exclusive H3K4me3-BDs (for instance *CD2*, *GATA3*, *BCL11B* or *CD28*) and myeloid identity genes within myeloid exclusive H3K4me3-BDs (for instance *TAL1*, *CEBPA*, *IL18* or *IL1B*). In comparison, H3K4me3-BDs that were shared between B cells, T cells and myeloid cells were associated with genes involved in basic cell functions such as RNA processing or epigenetic modifications and so they are likely reflecting the house-keeping background rather than haematopoiesis.

Total of 64, 74 and 206 H3K4me3-BDs were identified in MCL, CLL and MM that were not present in B cells. The high number of H3K4me3-BDs observed in MM can be due to the exclusion of H3K4me3-BDs from B cells which may not be representative of myeloma cells derived from plasma cells. Analysis of H3K4me3-BDs in B-cell malignancies for example identified T-cell markers such as *CD2* or *CD247*, proto-oncogenes *YES1*, *ETV1*, or *MYCN* or other cancer-associated genes such as *SDC1*, *WNT5A*, *SIX4* or *BCAR1*. The complete list of H3K4me3-BDs and their epigenomic specificity and gene annotation is provided in **Supplemental Table S4**.

We observed a significantly higher co-occurrence of H3K4me3-BD and super-enhancers compared to H3K4me3-BD and promoters (**Supplemental Fig. S4**). This enrichment increased with H3K4me3-BD size and was highly significant within a 100 kb proximity across all tested healthy and malignant cell types. Co-occurrence with super-enhancers within 100 kb was stronger in H3K4me3-BDs exclusive for B cells and myeloid cells (**Supplemental Fig. S4D,F**). T-cell exclusive H3K4me3-BDs showed broader enrichment with super-enhancers present more frequently within 2 Mb proximity (**Supplemental Fig.S4E**). This may be a result of a small number of H3K4me3-BDs, but also it could be a specific mechanism present in T cells, highlighting the presence of longer-range genomic interactions.

**Supplemental Tables**

**Supplemental Table S1: Selected BLUEPRINT chromatin states and their characterization.**

| **Original #[1]** | **Description** | **Histone marks high signal** | **Figure colour** |
|---|---|---|---|
| State 9 | Active canonical enhancer | H3K4me1 & H3K27ac | Orange |
| State 10 | Active non-canonical H3K4me3+ enhancer | H3K4me1 & H3K4me3 & H3K27ac | Green |
| State 11 | Active promoter | H3K4me3 & H3K27ac | Yellow |
| State 7 | Repressed Polycomb regulatory region | H3K4me1 & H3K4me3 & H3K27me3 | Light blue |

[1]Carrillo-de-Santa-Pau, Enrique, David Juan, Vera Pancaldi, Felipe Were, Ignacio Martin-Subero, Daniel Rico, Alfonso Valencia, and on behalf of The BLUEPRINT Consortium. 2017. “Automatic Identification of Informative Regions with Epigenomic Changes Associated to Hematopoiesis.” Nucleic Acids Research.

**Supplemental Table S2: Overview of BLUEPRINT ChIP-seq samples from healthy donors and patients with B-cell haematological malignancies.**

**Healthy donors**

| Cell lineage | Cell type | Cell type detail | Markers | Tissue | n |
|---|---|---|---|---|---|
| - | HSC | Mobilized primary cells | CD34+ | CB | 3 |
| Lymphoid | B cells | Naïve | $CD19^{+}CD27^{-}IgD^{+}$ | VB or CB | 6 |
| | | GC | $CD19^{+}CD20^{hi}CD38^{med}$ | T | 3 |
| | | Unswitched memory | $CD19^{+}CD27^{+}IgM^{+}IgD^{+}$ | VB | 1 |
| | | Class-switched memory | $CD19^{+}CD27^{+}IgA^{+}IgG^{+}$ or $CD19^{+}CD27^{-}IgD^{-}$ | VB | 3 |
| | Plasma cells | - | $CD20^{med}CD38^{high}$ | T | 2 |
| | T cells | Early cortical | $TCR\alpha\beta^{-}CD3^{-}CD4^{+}CD8^{+}$ | Th | 1# |
| | | Late cortical | $TCR\alpha\beta^{+}CD3^{+low}CD4^{+}CD8^{+}$ | Th | 1# |
| | | Helper, alphabeta | $CD3^{+}CD4^{+}CD45RA^{+}$ | VB or CB | 9 |
| | | Helper, alphabeta, central memory | $CD3^{+}CD4^{+}CD45RA^{-}CD62^{+}$ | VB | 1 |
| | | Cytotoxic, alphabeta | $CD3^{+}CD8^{+}CD45RA^{+}$ | VB or CB | 4 |
| | | Cytotoxic, alphabeta, effector memory | $CD3^{+}CD8^{+}CD45RA^{-}CD62L^{-}$ | VB | 1 |
| Myeloid | Neutrophils | Mature | $CD66b^{+}CD16^{+}$or NA | VB or CB | 13 |
| | | Band form | $CD11b^{+}CD16^{dim}$ | BM | 3 |
| | | Segmented | $CD11b^{+}CD16^{++}$ | BM | 2 |
| | Eosinophils | Mature | $CD66^{+}CD16^{-}$ | VB | 2 |
| | Monocytes | Classical | $CD14^{+}CD16^{-}$ | VB or CB | 5 |
| | Macrophages | Resting (M0) | * | VB or CB | 7 |
| | | Classically activated (M1) | * | VB or CB | 7 |
| | | Alternatively activated (M2) | * | VB or CB | 7 |
| | Erythroblasts | - | $CD36^{+}CD71^{+}CD235a^{+}$ | CB | 2 |
| | Megakaryocytes | - | $CD34^{-}CD41^{+}CD42^{+}$ | CB | 2 |

**Patients with B-cell haematological malignancies**

| Malignancy | Cell type | Markers | Tissue | n |
|---|---|---|---|---|
| CLL | Mature naïve (uCLL) or post GC B cells (mCLL) | - | VB | 7 |
| MCL | Mature naïve pre-GC B cells | - | VB | 5 |
| MM | Plasma cells | $CD138^{+}$ | BM | 4 |

Abbreviations: HSC – Haematopoietic stem cells, GC – Germinal centre, CB – Cord blood, VB – Venous blood, T – Tonsils, Th – Thymus, BM – Bone marrow, CLL – Chronic lymphocytic leukemia, uCLL – CLL with unmutated Ig variable region, mCLL – CLL with somatically mutated immunoglobulin variable region, MCL – Mantle cell lymphoma, MM – Multiple myeloma, *for details see BLUEPRINT methods. Data source: Carrillo-de-Santa-Pau E, Juan D, Pancaldi V, Were F, Martin-Subero I, Rico D, et al. Automatic identification of informative regions with epigenomic changes associated to hematopoiesis. Nucleic Acids Res. 2017;45(16):9244-59. #ChIP-seq reads from multiple donors merged, for details see Cieslak A, Charbonnier G, Tesio M, Mathieu EL, Belhocine M, Touzart A, et al. Blueprint of human thymopoiesis reveals molecular mechanisms of stage-specific TCR enhancer activation. J Exp Med. 2020;217(9).

**Supplemental Table S3: Characteristics of cell line derived from B-cell and T-cell haematological malignancies, data availability and sources.**

| Cell line | Malig-nancy | Cell type origin | Sex | Chromosomal translocation | Data source[#] | | |
|---|---|---|---|---|---|---|---|
| | | | | | ChIP-seq | DNase I HS & RNA-seq | TS |
| JVM-2 | MCL | Mature naïve pre-GC B cells | F | t(11;14)(q13;q32) *CCND1-IGH* | BLUEPRINT | | |
| Z-138 | MCL | Mature naïve pre-GC B cells | M | t(8;11;14)(q24;q13;q32) *MYC-CCND1-IGH* | BLUEPRINT | BLUEPRINT | |
| BL-2 | BL | GC B cells | M | t(8;22)(q24;q11.2) *MYC-IGL* | BLUEPRINT | | |
| DG-75 | BL | GC B cells | M | t(8;14)(q24;q32) *MYC-IGH* | BLUEPRINT | BLUEPRINT | |
| KARPAS-422 | DLBCL | GC B cells | F | t(14;18)(q32;q21) *IGH-BCL2* | BLUEPRINT | BLUEPRINT | |
| SU-DHL-5 | DLBCL | GC B cells | F | none | BLUEPRINT | BLUEPRINT | |
| U266 | MM | Plasma cells | M | t(11;14)(q13;q32) *CCND1-IGH* | BLUEPRINT | BLUEPRINT | MM |
| KMS11 | MM | Plasma cells | F | t(4;8;14;16)(p16;q24;q32;q23) *NSD2-MYC-IGH-MAF* | MM | | MM |
| MM1S | MM | Plasma cells | F | t(8;14;16)(q24;q32;q23) *MYC-IGH-MAF* | MM | | MM |
| KOPT-K1 | T-ALL | Immature T cells | M | t(11;14)(p13;q11) *LMO2-TRA/TRD* | GSE54379 | | |
| DND-41 | T-ALL | Immature T cells | M | t(5;14)(q35;q32) *TLX3-BCL11B* | GSE54379 | | |
| Jurkat | T-ALL | Immature T cells | M | none* | GSE65687 | | |
| PEER | T-ALL | Immature T cells | F | none* | T-ALL | | |

Abbreviations: GC – Germinal center, DNase I HS – DNase I hypersensitivity, TS – Targeted DNA sequencing, MCL – Mantle cell lymphoma, BL – Burkitt lymphoma, DLBCL – Diffuse large B-cell lymphoma, MM – Multiple myeloma; T-ALL – T acute lymphoblastic leukemia. *Jurkat cell line has a 12 bp insertion upstream of the *TAL1* gene that generates a *de novo* super-enhancer. This insertion was introduced in PEER cell line using CRISPR-Cas9.

[#]Notes for data source:
BLUEPRINT: Carrillo-de-Santa-Pau E, Juan D, Pancaldi V, Were F, Martin-Subero I, Rico D, et al. Automatic identification of informative regions with epigenomic changes associated to hematopoiesis. Nucleic Acids Res. 2017;45(16):9244-59.
MM: ChIP-seq and DNA sequencing data of U266, KMS11 and MM1S provided by Brian Walker.
GSE54379: https://www.ncbi.nlm.nih.gov/geo/query/acc.cgi?acc=GSE54379.
GSE65687: https://www.ncbi.nlm.nih.gov/geo/query/acc.cgi?acc=GSE65687.
T-ALL: ChIP-seq of wild type and edited PEER cell line data provided by Salvatore Spicuglia.

**Supplemental Table S4: List of 59,097 H3K4me3-BDs detected by genome-wide analysis.** Abbreviations: HSC – Haematopoietic stem cells, B – B cells, T – T cells, M – Myeloid cells, MCL – Mantle cell lymphoma, CLL – Chronic lymphocytic leukemia, MM – Multiple myeloma. Overlapped H3K4me3-BDs can be recognized by the same labelling within "PART" column.

*Supplemental Table S4 is attached as a separated file.*

**Supplemental Table S5: Characteristics of seven patient derived xenograft multiple myeloma samples.**

| **ID** | **Malig-nancy** | **Cell type origin** | **Sex** | **Chromosomal translocation** | **ChIP-seq** | **RNA-seq** | **WGS** |
|---|---|---|---|---|---|---|---|
| **P1** | MM | Plasma cells | F | t(14;16)(q32;q23) *IGH-MAF* | yes | yes | yes |
| **P2** | MM | Plasma cells | F | t(14;16)(q32;q23) *IGH-MAF* | yes | - | yes |
| **P3** | MM | Plasma cells | F | t(4;14)(p16;q32) *NSD2-IGH* | yes | yes | yes |
| **P4** | MM | Plasma cells | M | Complex translocation involving *NSD2* and *IGH* | yes | yes | yes |
| **P5** | MM | Plasma cells | F | t(4;8)(q28;q24) *PCHD10-MYC* | yes | yes | yes |
| **P6** | MM | Plasma cells | M | none | yes | yes | yes |
| **P7** | MM | Plasma cells | M | none | yes | - | yes |

Abbreviations: MM – Multiple myeloma, WGS – Whole genome sequencing.

### Supplemental Figures

**Supplemental Figure S1: Immunoglobulin structure, genes and physiological genomic rearrangements.** Abbreviations: C – Constant region, J – Joining region, D – Diversity region, V – Variable region, RAG – Recombination-activating gene, AID – Activation-induced cytidine deaminase.

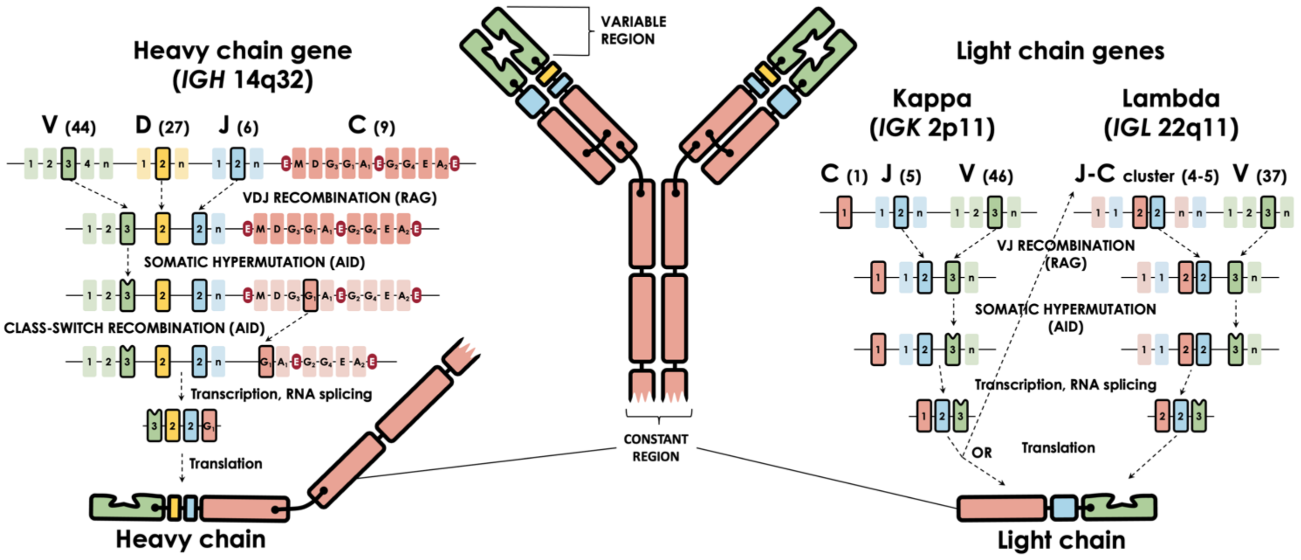

**Supplemental Figure S2: Overview of H3K4me3-BDs detected at genome-wide level**. Classification of H3K4me3-BD epigenomic specificity across healthy haematopoiesis and B-cell malignancies and proportions of each category within each group is shown. Abbreviations: H – Haematopoietic stem cells, B – B cells, T – T cells, M – Myeloid cells, MCL – Mantle cell lymphoma, CLL – Chronic lymphocytic leukemia, MM – Multiple myeloma.

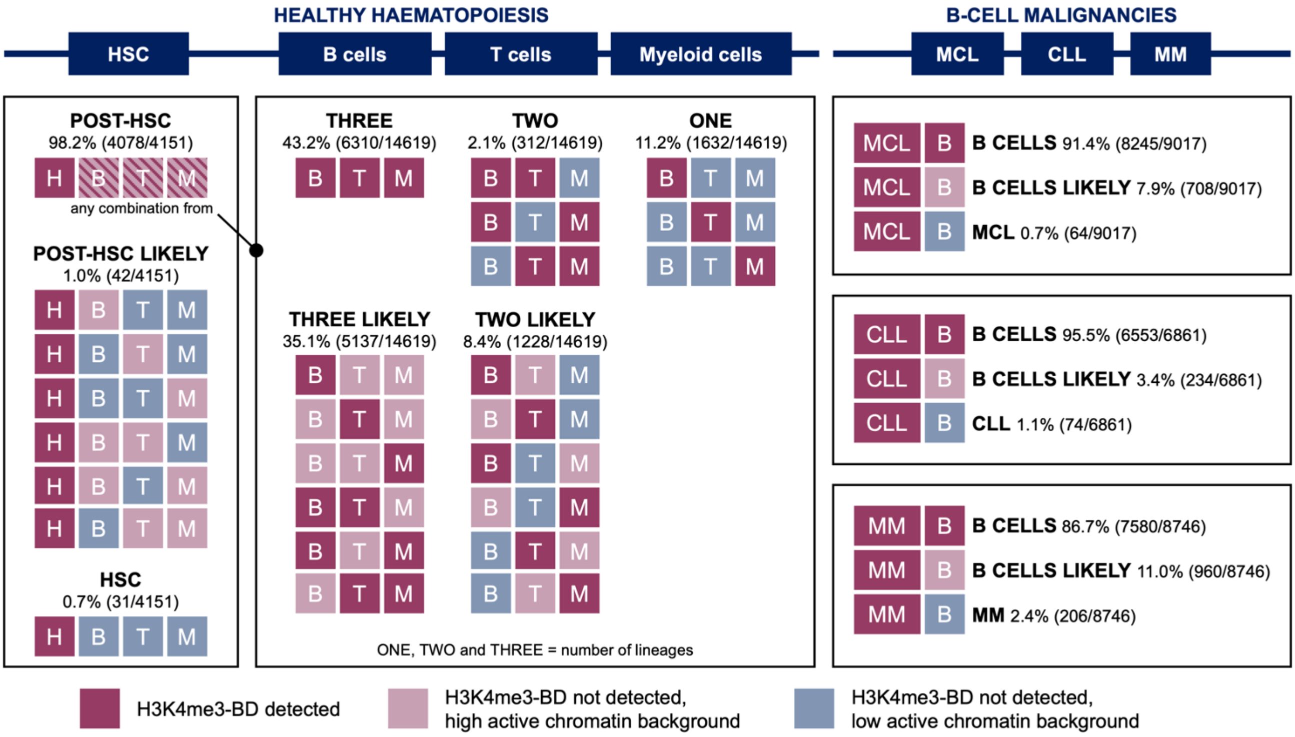

**Supplemental Figure S3: Genes annotated to H3K4me3-BD enrichment analysis for biological processes.** Selected categories of H3K4me3-BDs from **Supplemental Fig. S2** are shown as follow: H3K4me3-BDs exclusive for B cells **(A)**, T cells **(B)** and myeloid cells **(C)**, H3K4me3-BDs detected in all three lineages (B cells, T cells and myeloid cells) **(D)**, H3K4me3-BDs exclusive for haematopoietic stem cells **(E)** and for multiple myeloma **(F)**. First top 10 statistically significant biological processes are displayed. No significant enrichment was found in mantle cell lymphoma and chronic lymphocytic leukemia. Abbreviations: B – B cells, T – T cells, M – Myeloid cells, H – Haematopoietic stem cells, MM – Multiple myeloma.

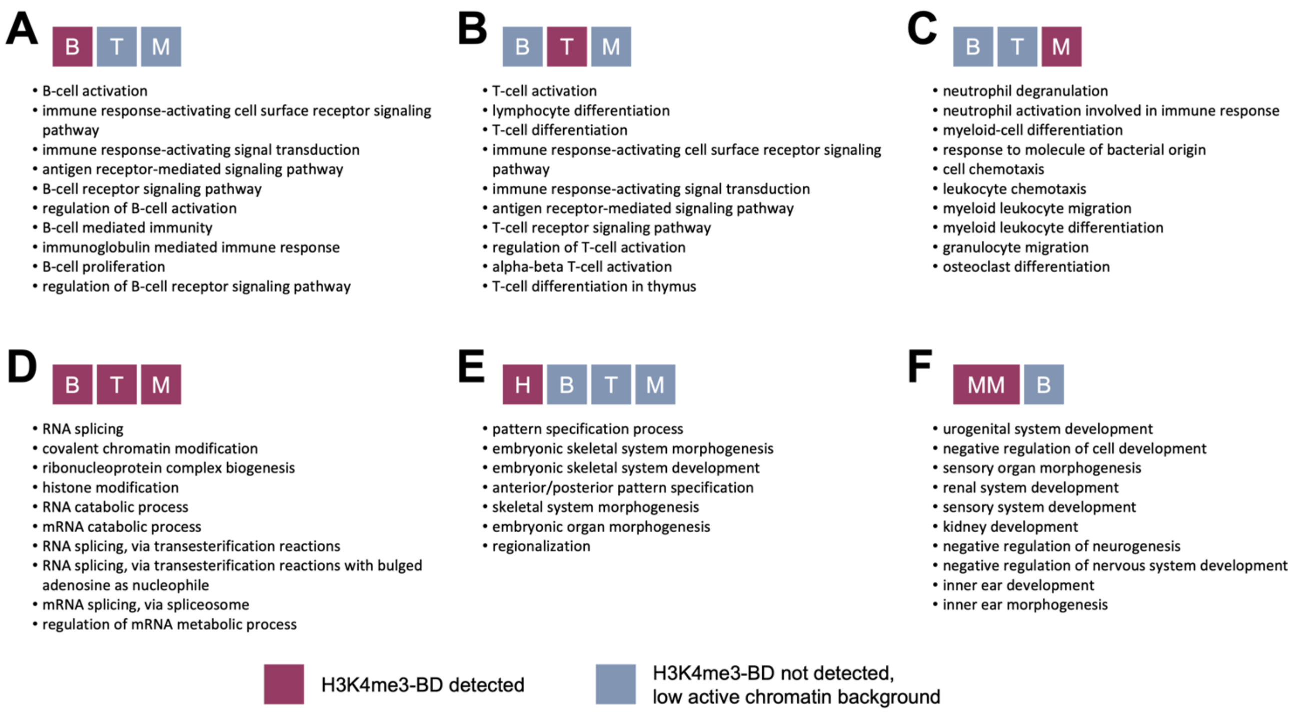

**Supplemental Figure S4: Proximity of super-enhancers to H3K4me3-BDs or small promoters.** Each chart shows a fraction of H3K4me3-BDs (red dots) and promoters (blue dots) that have super-enhancer(s) within 100 kb distance windows in B cells **(A, D)**, T cells **(B, E)**, myeloid cells **(C, F)**. Upper part **(A–C)** includes all H3K4me3-BDs detected in the given cell type and the lower part **(D–F)** shows only those H3K4me3-BDs recognized as cell type exclusive. Data for each chart are divided into two H3K4me3-BDs size intervals: from 2 kb to smaller than 5 kb (marked as <5 kb, on left) and 5 kb and bigger (marked ≥5 kb, on right). H3K4me3-BDs and promoter fractions were compared by Fisher's exact test. Statistically significant levels are as follows: ***P<0.001, **P<0.01 and *P<0.05. Abbreviations: B – B cells, T – T cells, M – Myeloid cells.

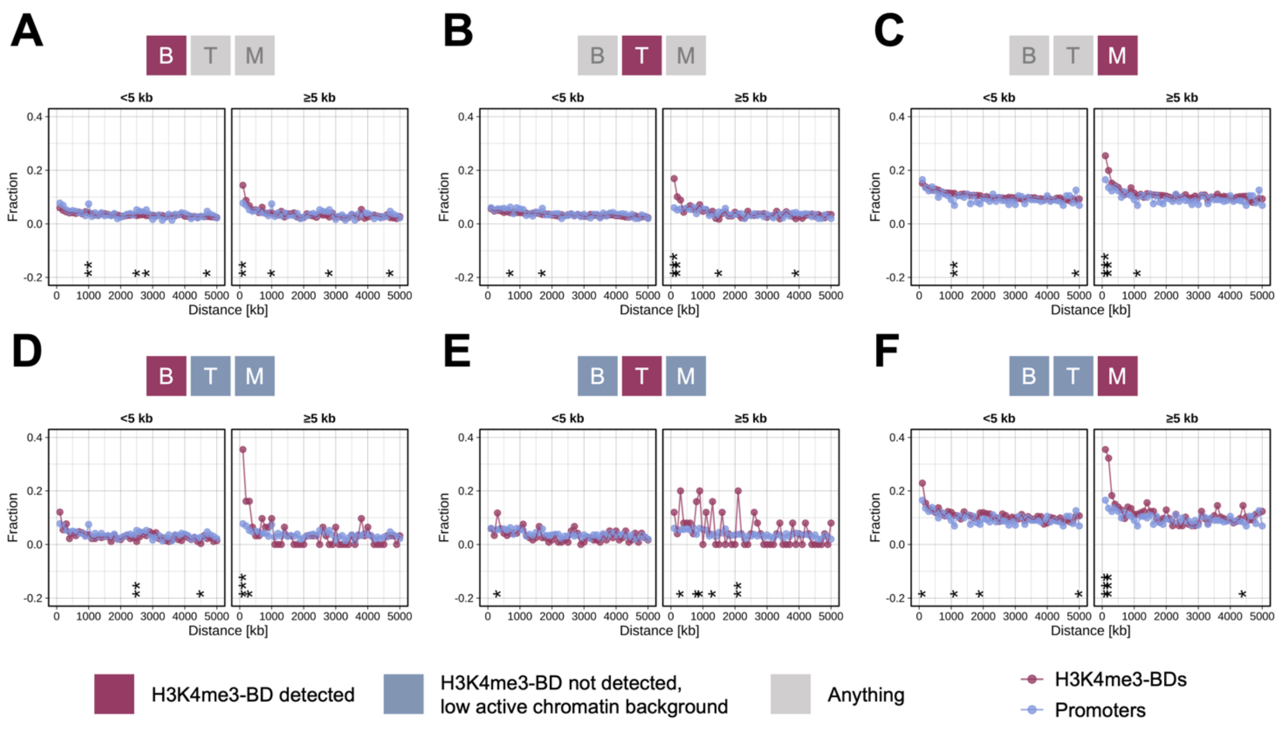

**Supplemental Figure S5: Chromatin landscape of the human *IGH* locus (14q32) in five cell lines derived from B-cell haematological malignancies.** Upper line of each cell line (**Supplemental Table S3**) represents the selected ChIP-seq chromatin states (**Supplemental Table S1**) and the lower line shows DNase I hypersensitivity sites for the non-variable region of the *IGH* locus. Vertical black lines in sample U266 mark breakpoint positions for the region, which includes the Eα1 super-enhancer that inserts ~12 kb upstream of *CCND1* (11q13). (**Fig. 3**). Abbreviations: BD – Broad domain.

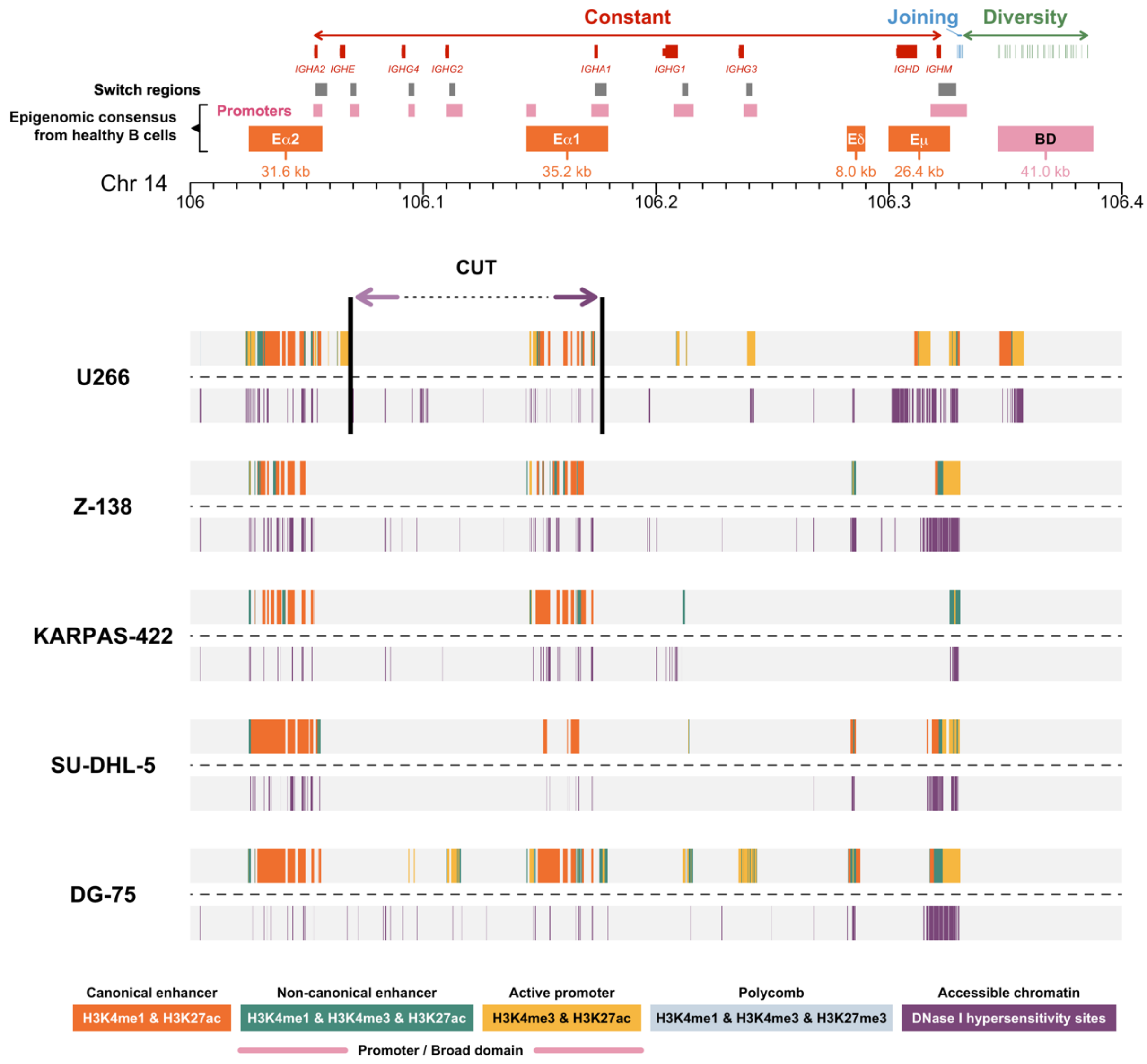

**Supplemental Figure S6: Chromatin landscape of the *MYEOV* locus (11q13) in healthy human B cells and five cell lines derived from B-cell haematological malignancies.** Selected ChIP-seq chromatin states (**Supplemental Table S1**) of *MYEOV* gene in BLUEPRINT healthy B cells are shown in upper part. Upper line of each BLUEPRINT cell line in lower part represents the selected ChIP-seq chromatin states and the lower line shows DNase I hypersensitivity sites for the *MYEOV* locus. Numbers in coloured squares (red denotes high expression and blue low expression) show *MYEOV* expression detected using RNA-seq in Fragments Per Kilobase of transcript per Million mapped reads (FPKM). Abbreviations: GC – Germinal centre.

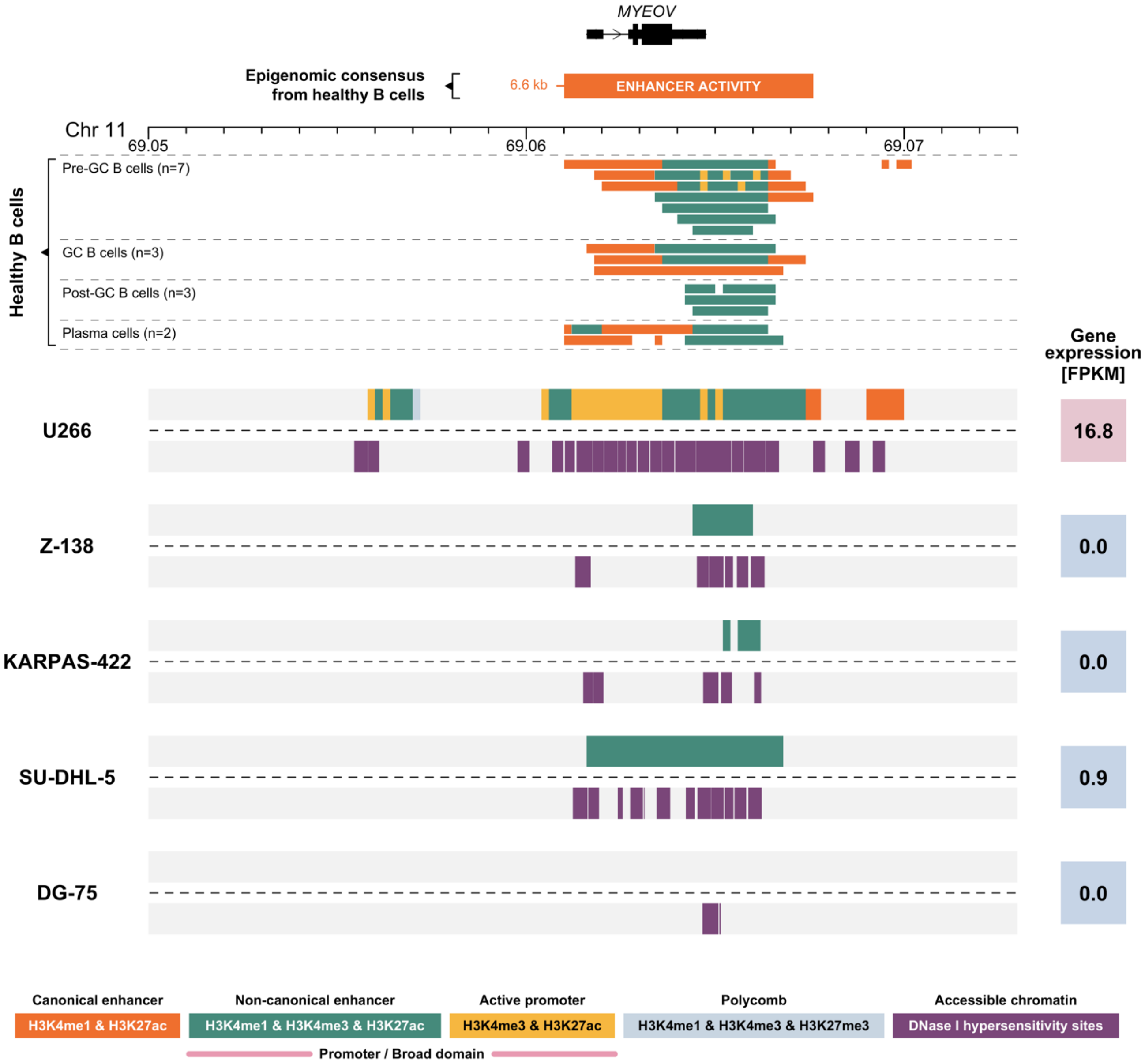

**Supplemental Figure S7: Chromatin landscape of the *MYC* (8q24) and *FGFR3/NSD2* (4p16) loci in healthy human B cells and multiple myeloma samples.** Upper panels show selected ChIP-seq chromatin states (**Supplemental Table S1**) of *MYC* **(A)** and *FGFR3/NSD2* **(B)** loci in BLUEPRINT healthy B cells. Each line of lower panel represents the ChIP-seq chromatin states for myeloma cell lines KMS11 and MM1S and seven multiple myeloma patients (P1-P7, patient derived xenograft material) for the *MYC* **(A)** and the *FGFR3/NSD2* **(B)** loci. Numbers in coloured squares (red denotes high expression and blue low expression; the two shades of blue for *MYC* expression denote different levels of low expression determined in our previous study[1]) show gene expression detected by RNA-seq and displayed as $Log_2$ normalised counts ($Log_2$ NC). *sample contains chromosomal translocation involving the displayed region, described in detail in **Supplemental Tables S3 and S5**. Detailed picture of individual histone marks of the *MYC* and *FGFR3/NSD2* loci in multiple myeloma samples is given in **Supplemental Fig. S8**. Abbreviations: GC – Germinal centre.

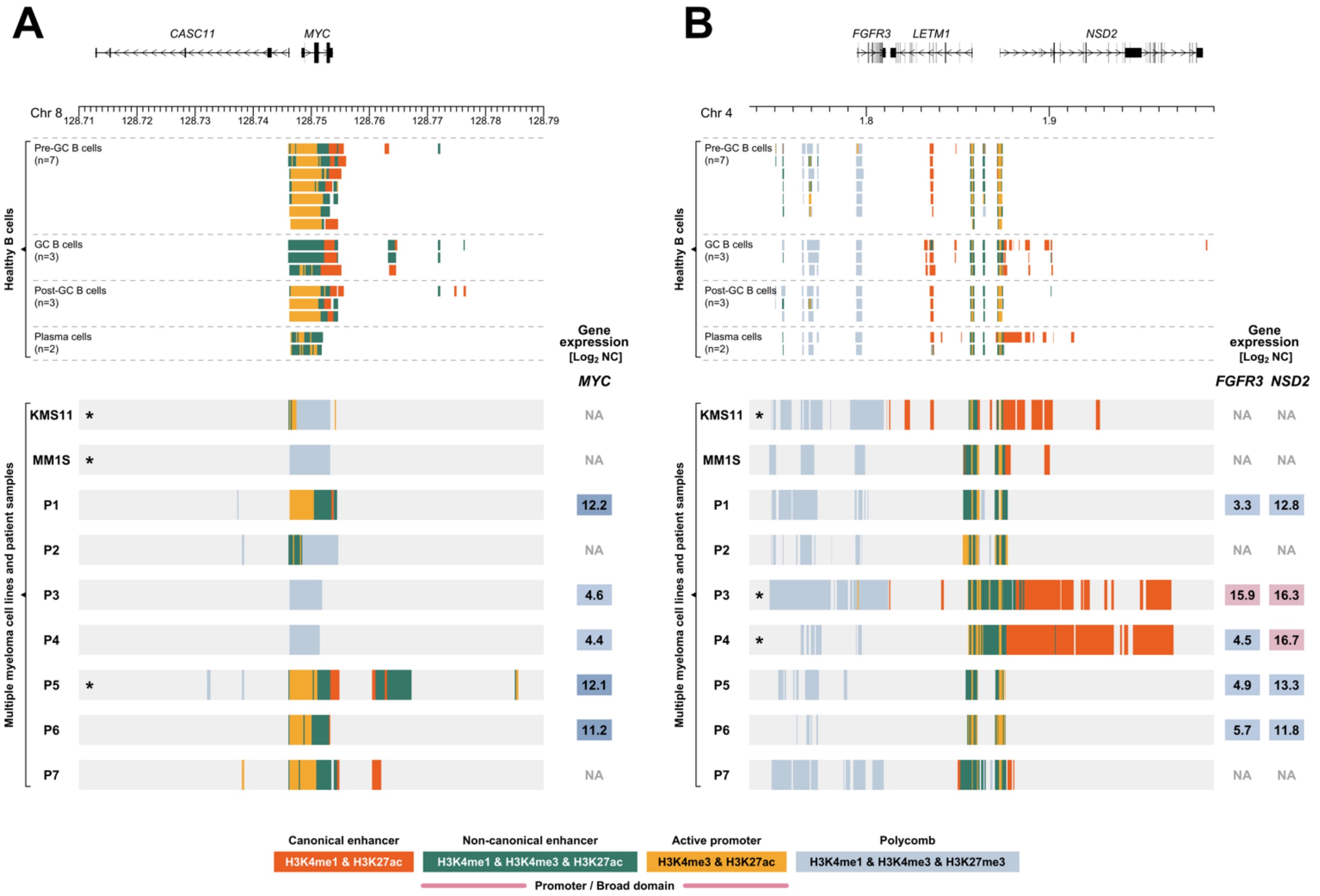

[1]Mikulasova, Aneta, Cody Ashby, Ruslana G. Tytarenko, Pingping Qu, Adam Rosenthal, Judith A. Dent, Katie R. Ryan, et al. 2019. "Microhomology-Mediated End Joining Drives Complex Rearrangements and over Expression of *MYC* and *PVT1* in Multiple Myeloma." Haematologica.

**Supplemental Figure S8: Detailed chromatin landscape of *MYC* (8q24) and *FGFR3/NSD2* (4p16) loci in multiple myeloma samples.** Each line represents ChIP-seq results for myeloma cell lines KMS11 and MM1S and seven multiple myeloma patients P1-P7, patient derived xenograft material) for the *MYC* **(A)** and the *FGFR3/NSD2* **(B)** loci. First line of each sample shows the selected ChIP-seq chromatin states (**Supplemental Table S1**) and other lines display peaks of individual histone marks. Numbers in coloured squares (red denotes high expression and blue low expression; the two shades of blue for *MYC* expression denote different levels of low expression determined in our previous study[1]) show gene expression detected by RNA-seq and displayed as $Log_2$ normalised counts ($Log_2$ NC). *sample contains chromosomal translocation involving the displayed region, described in detail in **Supplemental Tables S3 and S5**.

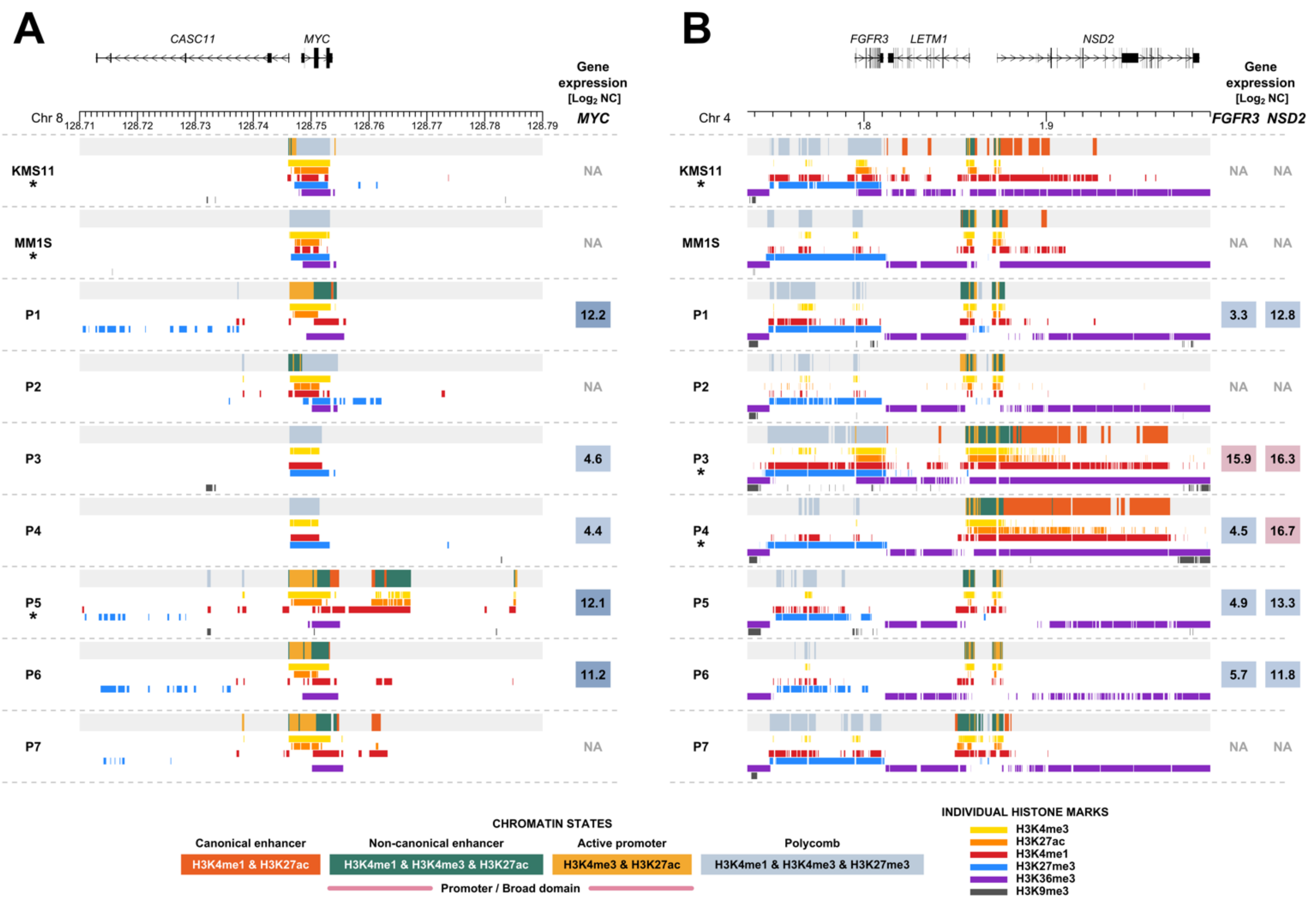

[1]Mikulasova Aneta, Cody Ashby, Ruslana G. Tytarenko, Pingping Qu, Adam Rosenthal, Judith A. Dent, Katie R. Ryan, et al. 2019. “Microhomology-Mediated End Joining Drives Complex Rearrangements and over Expression of *MYC* and *PVT1* in Multiple Myeloma.” Haematologica.

**Footnote:**

The H3K4me3-BD over *MYC* was missed by the chromatin state model in cell lines KMS11 and MM1S. This pattern was also observed in patient P2 despite no evidence of a translocation event. This observation remains unexplained. Additionally, a different epigenomic pattern over *MYC* was found in patient P5 with t(4;8). This difference can be explained by the presence of copy-number neutral loss of heterozygosity at 8q involving *MYC* locus and involvement of both alleles in *MYC* translocation, confirmed by whole genome sequencing alignments (data not shown). KMS11, P3 and P4 samples with a translocation involving the *FGFR3*/*NSD2* locus show a similar epigenomic pattern over the two genes, excluding *FGFR3* for patient P4. This difference is explained by the presence of a complex translocation in P4 sample involving monoallelic deletion of *FGFR3* gene, confirmed by in-depth analysis of whole genome sequencing (data not shown). KMS11 expresses high levels of *FGFR3* and *NSD2* (559 and 49 FKPM, respectively).

**Supplemental Figure S9: Genomic and epigenomic architecture of the *TRA/TRD* locus (14q11) in healthy and malignant human haematopoietic cells.** Each panel represents collapsed cell-type specific signals of ChIP-seq chromatin states included in this study (**Supplemental Table S1**). Abbreviations: HSC – Haematopoietic stem cells, GC – Germinal centre, CLL – Chronic lymphocytic leukemia, MCL – Mantle cell lymphoma, DLBCL – Diffuse large B-cell lymphoma, BL – Burkitt lymphoma, MM – Multiple myeloma.

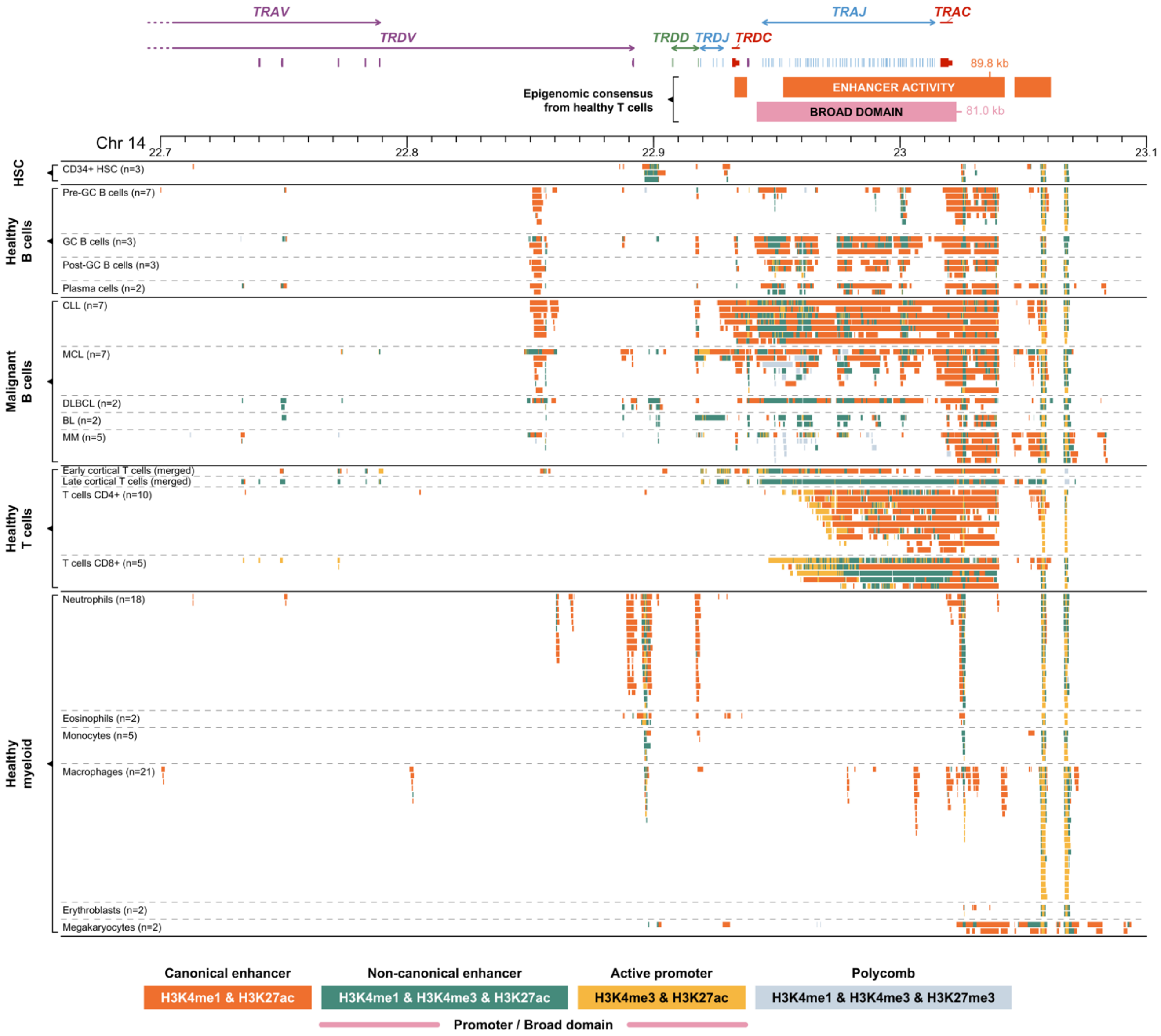

**Supplemental Figure S10: Genomic and epigenomic architecture of the *BCL11B* locus (14q32) in healthy and malignant human haematopoietic cells.** Each panel represents collapsed cell-type specific signals of ChIP-seq chromatin states included in this study (**Supplemental Table S1**). Areas with orange background mark regions showed in detail at **Supplemental Fig. S11**. Abbreviations: HSC – Haematopoietic stem cells, GC – Germinal centre, CLL – Chronic lymphocytic leukemia, MCL – Mantle cell lymphoma, DLBCL – Diffuse large B-cell lymphoma, BL – Burkitt lymphoma, MM – Multiple myeloma.

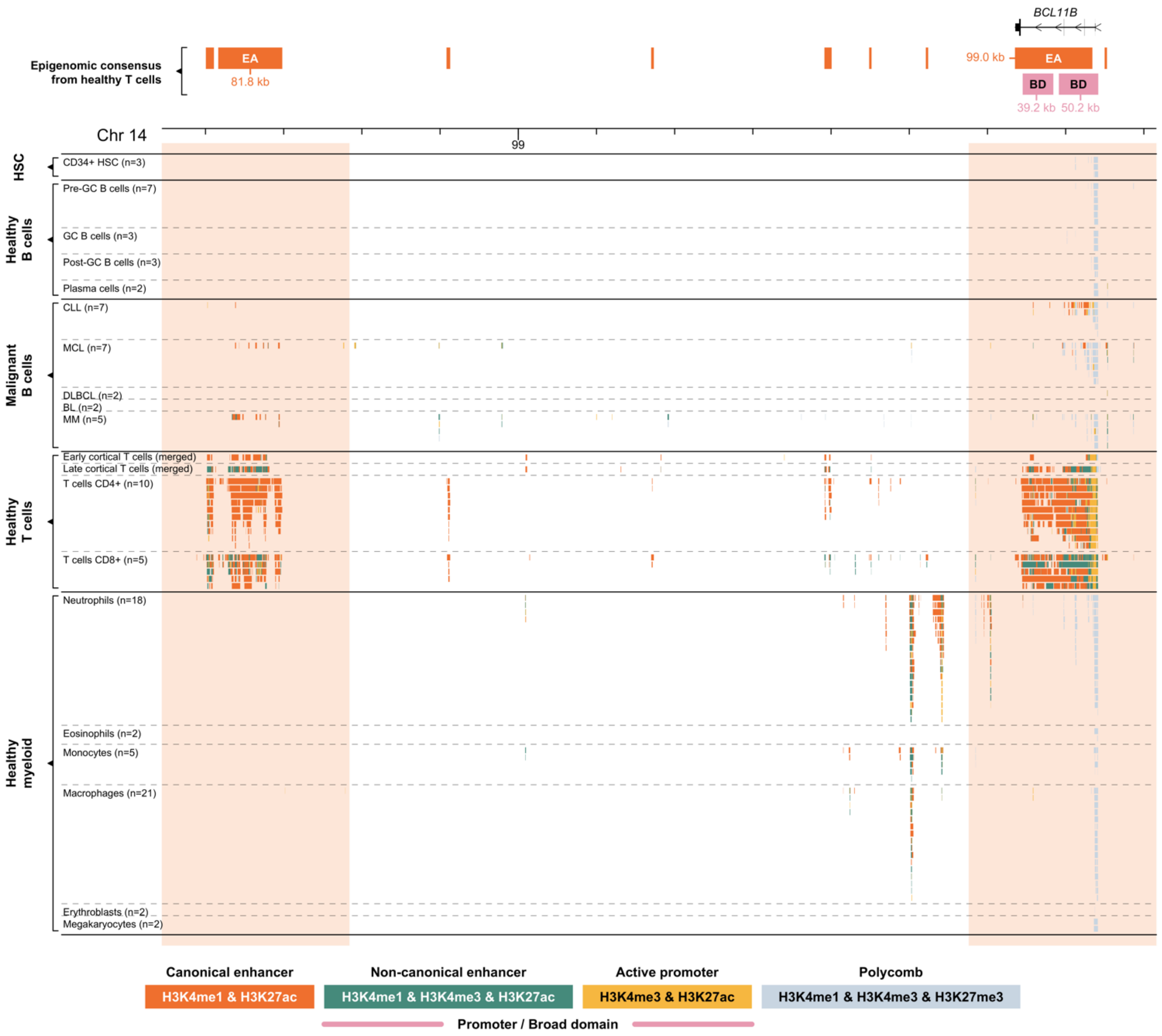

**Supplemental Figure S11: Genomic and epigenomic architecture of the *BCL11B* locus (14q32) in healthy and malignant human haematopoietic cells, detailed image.** Two regions marked by orange background in **Supplemental Fig. S10** are shown: intergenic super-enhancer activity in ~1 Mb proximity to *BCL11B* gene **(A)**, and *BCL11B* gene **(B)**. Each panel represents collapsed cell-type specific signals of ChIP-seq chromatin states included in this study (**Supplemental Table S1**). Abbreviations: HSC – Haematopoietic stem cells, GC – Germinal centre, CLL – Chronic lymphocytic leukemia, MCL – Mantle cell lymphoma, DLBCL – Diffuse large B-cell lymphoma, BL – Burkitt lymphoma, MM – Multiple myeloma.

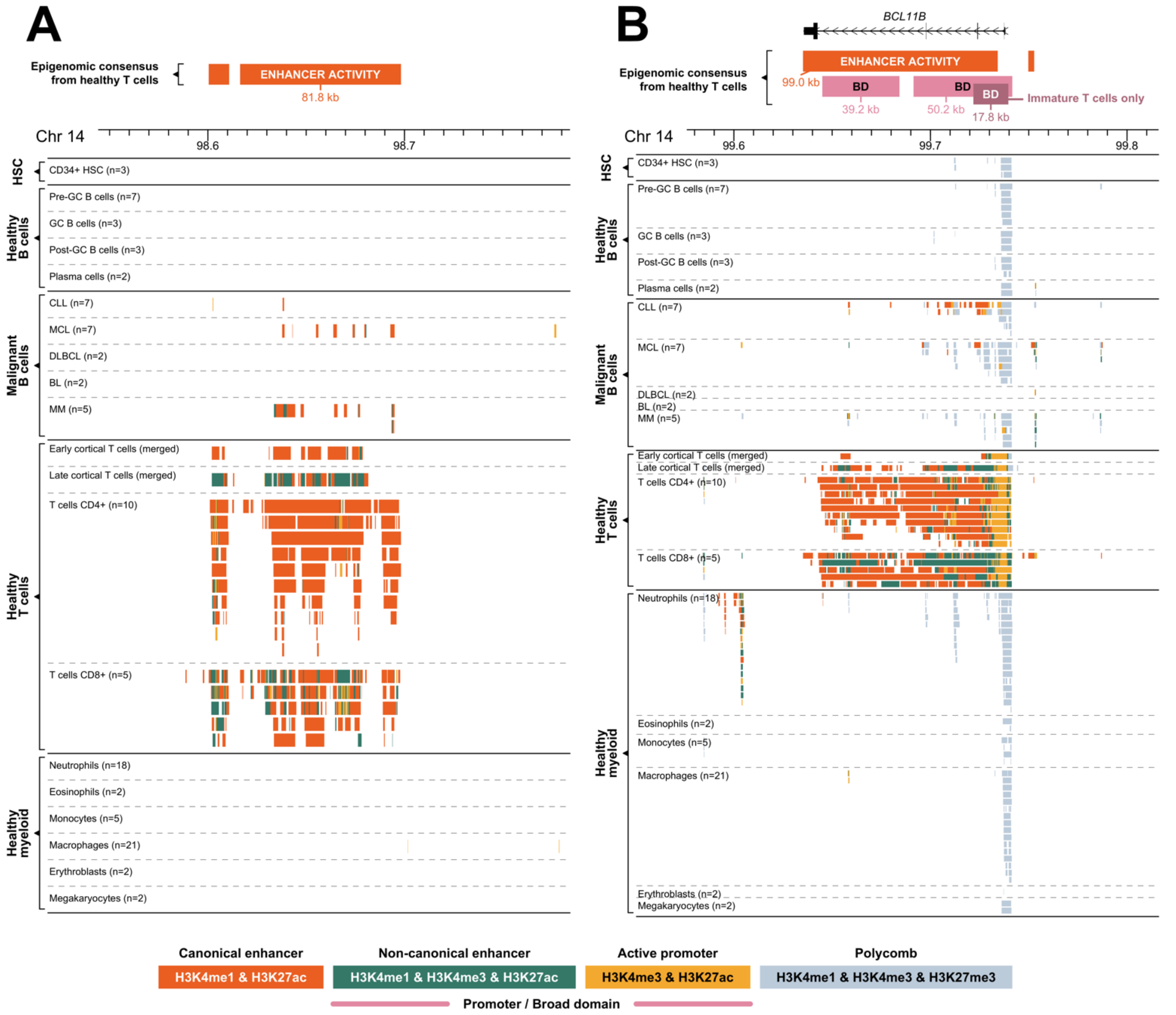

**Supplemental Figure S12: Genomic and epigenomic architecture of the *TAL1* locus (1p33) in healthy and malignant human haematopoietic cells.** Each panel represents collapsed cell-type specific signals of ChIP-seq chromatin states included in this study (**Supplemental Table S1**). Abbreviations: HSC – Haematopoietic stem cells, GC – Germinal centre, CLL – Chronic lymphocytic leukemia, MCL – Mantle cell lymphoma, DLBCL – Diffuse large B-cell lymphoma, BL – Burkitt lymphoma, MM – Multiple myeloma.

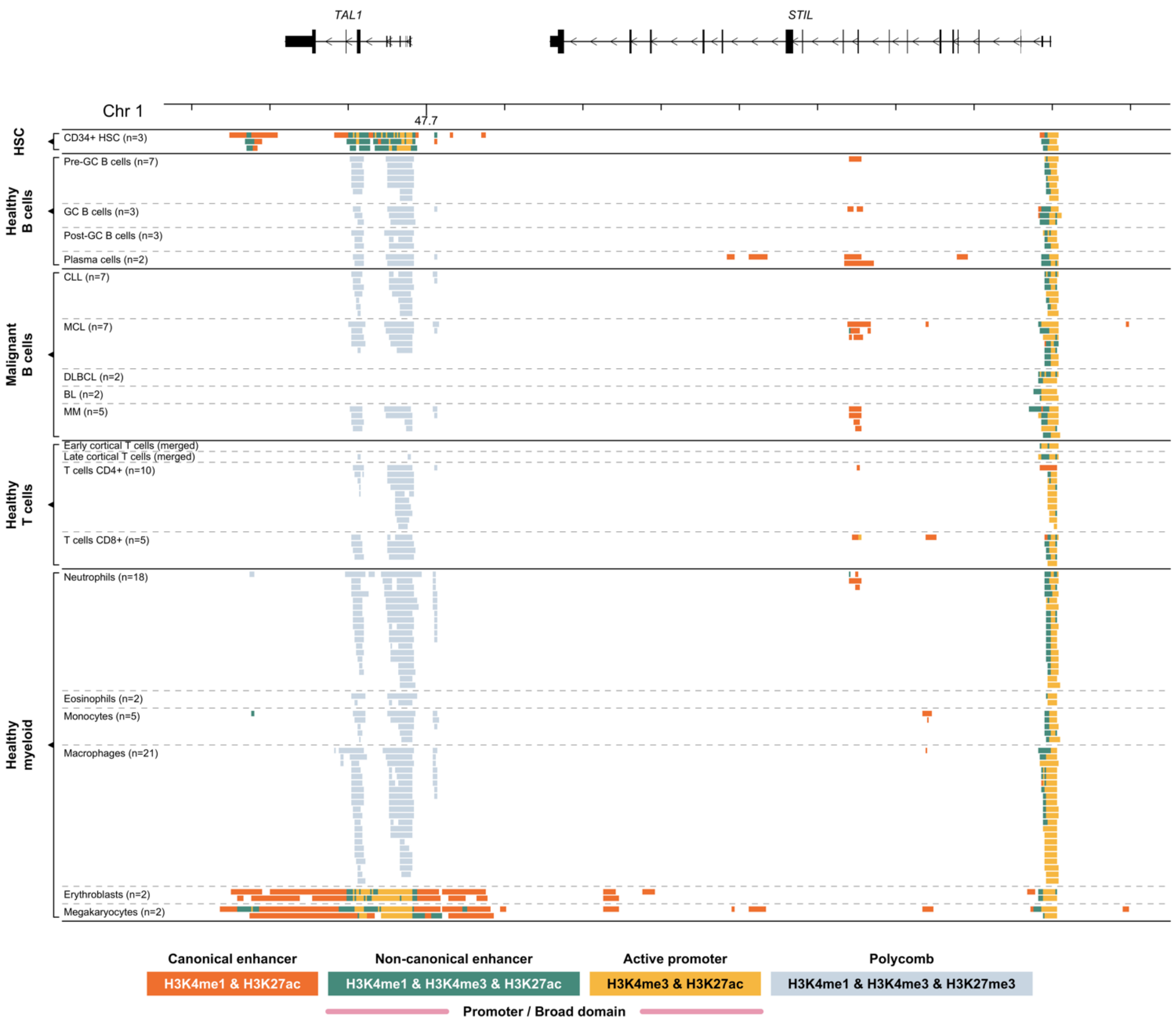

**Supplemental Figure S13: H3K4me3 and H3K27ac histone marks signal at *TAL1* locus in PEER cell line.** 12bp DNA insertion known from Jurkat cell line upstream of *TAL1* gene was introduced in PEER cell line using CRISPR-Cas9. Normalized read count is shown for wild-type PEER in black and for edited PEER in red.

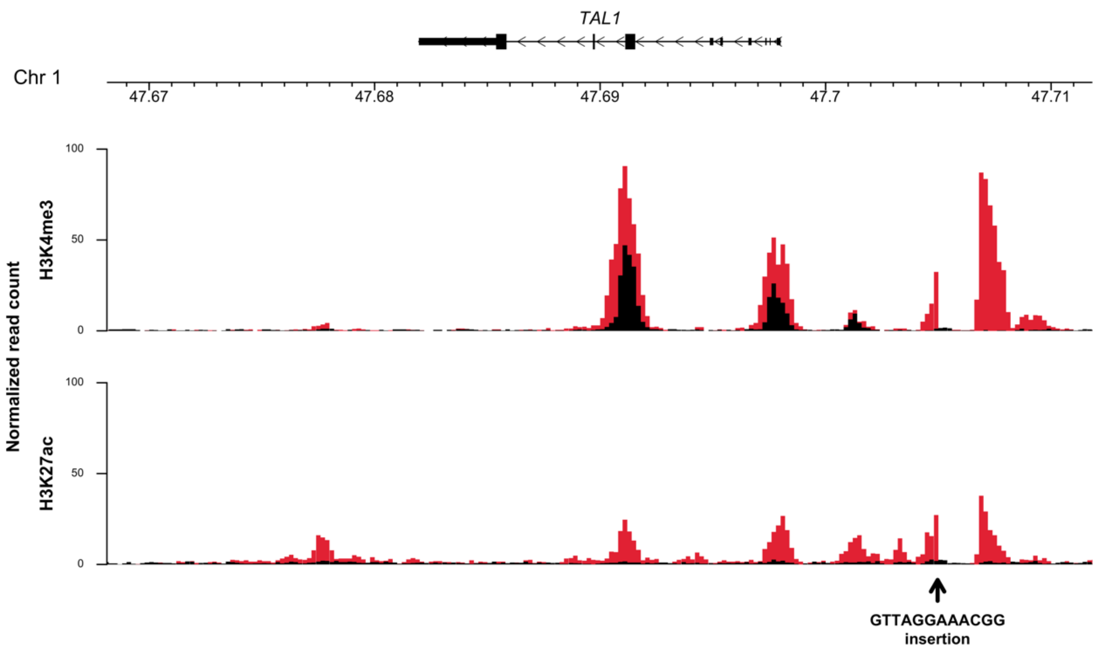
