## Supplementary material for "Epigenomic translocation of H3K4me3 broad domains over oncogenes following hijacking of super-enhancers": Supplmental Figures S1-13

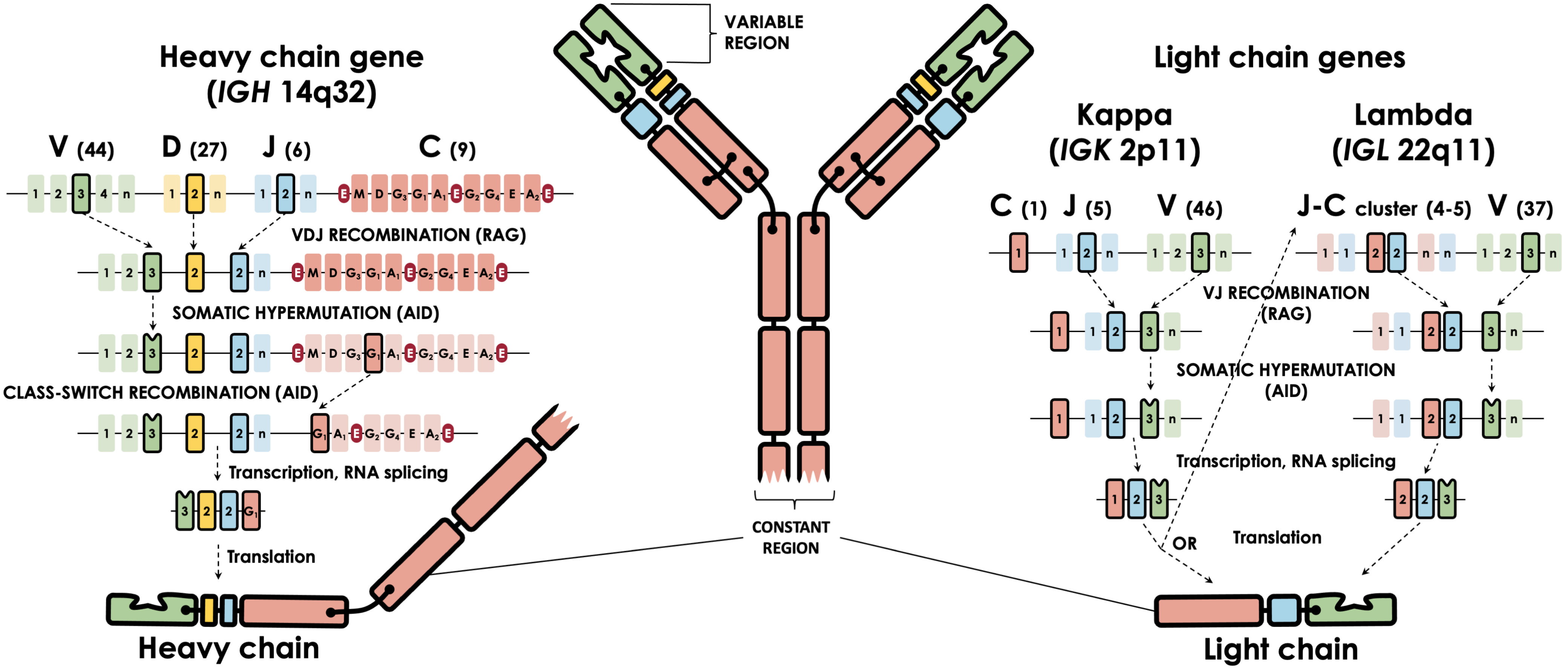
Heavy chain gene
(IGH 14q32)
V (44)
D (27)
J (6)
C (9)
VDJ RECOMBINATION (RAG)
SOMATIC HYPERMUTATION (AID)
CLASS-SWITCH RECOMBINATION (AID)
Transcription, RNA splicing
Translation
Heavy chain
VARIABLE REGION
CONSTANT REGION
Light chain genes
Kappa
(IGK 2p11)
Lambda
(IGL 22q11)
C (1)
J (5)
V (46)
J-C cluster (4-5)
V (37)
VJ RECOMBINATION (RAG)
SOMATIC HYPERMUTATION (AID)
Transcription, RNA splicing
OR
Translation
Light chain

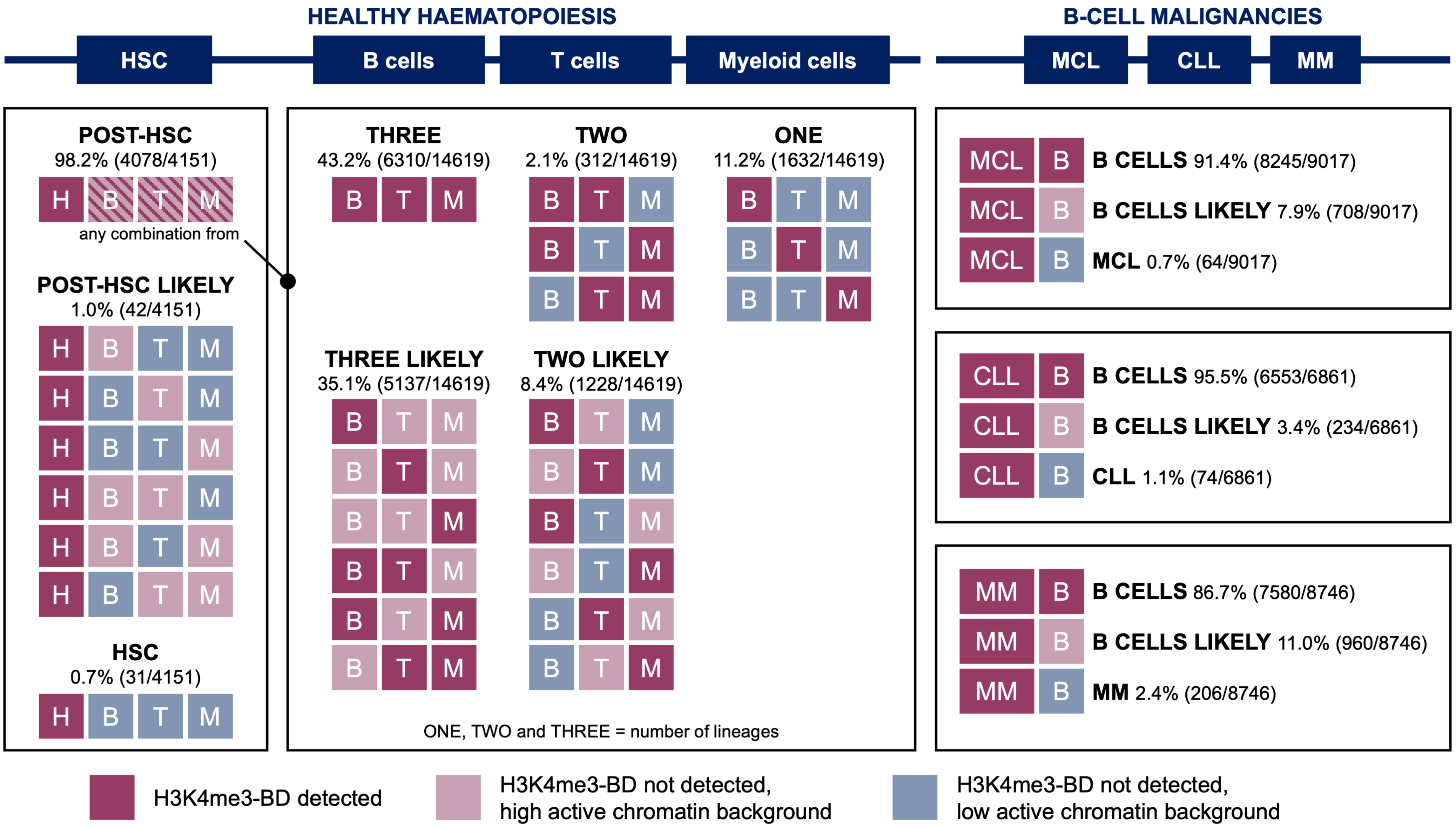

HEALTHY HAEMATOPOIESIS
B-CELL MALIGNANCIES
HSC
B cells
T cells
Myeloid cells
MCL
CLL
MM
POST-HSC
98.2% (4078/4151)
any combination from
POST-HSC LIKELY
1.0% (42/4151)
HSC
0.7% (31/4151)
THREE
43.2% (6310/14619)
TWO
2.1% (312/14619)
ONE
11.2% (1632/14619)
THREE LIKELY
35.1% (5137/14619)
TWO LIKELY
8.4% (1228/14619)
ONE, TWO and THREE = number of lineages
B CELLS 91.4% (8245/9017)
B CELLS LIKELY 7.9% (708/9017)
MCL 0.7% (64/9017)
B CELLS 95.5% (6553/6861)
B CELLS LIKELY 3.4% (234/6861)
CLL 1.1% (74/6861)
B CELLS 86.7% (7580/8746)
B CELLS LIKELY 11.0% (960/8746)
MM 2.4% (206/8746)
H3K4me3-BD detected
H3K4me3-BD not detected, high active chromatin background
H3K4me3-BD not detected, low active chromatin background

**A** B T M

- B-cell activation
- immune response-activating cell surface receptor signaling pathway
- immune response-activating signal transduction
- antigen receptor-mediated signaling pathway
- B-cell receptor signaling pathway
- regulation of B-cell activation
- B-cell mediated immunity
- immunoglobulin mediated immune response
- B-cell proliferation
- regulation of B-cell receptor signaling pathway

**B** B T M

- T-cell activation
- lymphocyte differentiation
- T-cell differentiation
- immune response-activating cell surface receptor signaling pathway
- immune response-activating signal transduction
- antigen receptor-mediated signaling pathway
- T-cell receptor signaling pathway
- regulation of T-cell activation
- alpha-beta T-cell activation
- T-cell differentiation in thymus

**C** B T M

- neutrophil degranulation
- neutrophil activation involved in immune response
- myeloid-cell differentiation
- response to molecule of bacterial origin
- cell chemotaxis
- leukocyte chemotaxis
- myeloid leukocyte migration
- myeloid leukocyte differentiation
- granulocyte migration
- osteoclast differentiation

**D** B T M

- RNA splicing
- covalent chromatin modification
- ribonucleoprotein complex biogenesis
- histone modification
- RNA catabolic process
- mRNA catabolic process
- RNA splicing, via transesterification reactions
- RNA splicing, via transesterification reactions with bulged adenosine as nucleophile
- mRNA splicing, via spliceosome
- regulation of mRNA metabolic process

**E** H B T M

- pattern specification process
- embryonic skeletal system morphogenesis
- embryonic skeletal system development
- anterior/posterior pattern specification
- skeletal system morphogenesis
- embryonic organ morphogenesis
- regionalization

**F** MM B

- urogenital system development
- negative regulation of cell development
- sensory organ morphogenesis
- renal system development
- sensory system development
- kidney development
- negative regulation of neurogenesis
- negative regulation of nervous system development
- inner ear development
- inner ear morphogenesis

H3K4me3-BD detected

H3K4me3-BD not detected, low active chromatin background

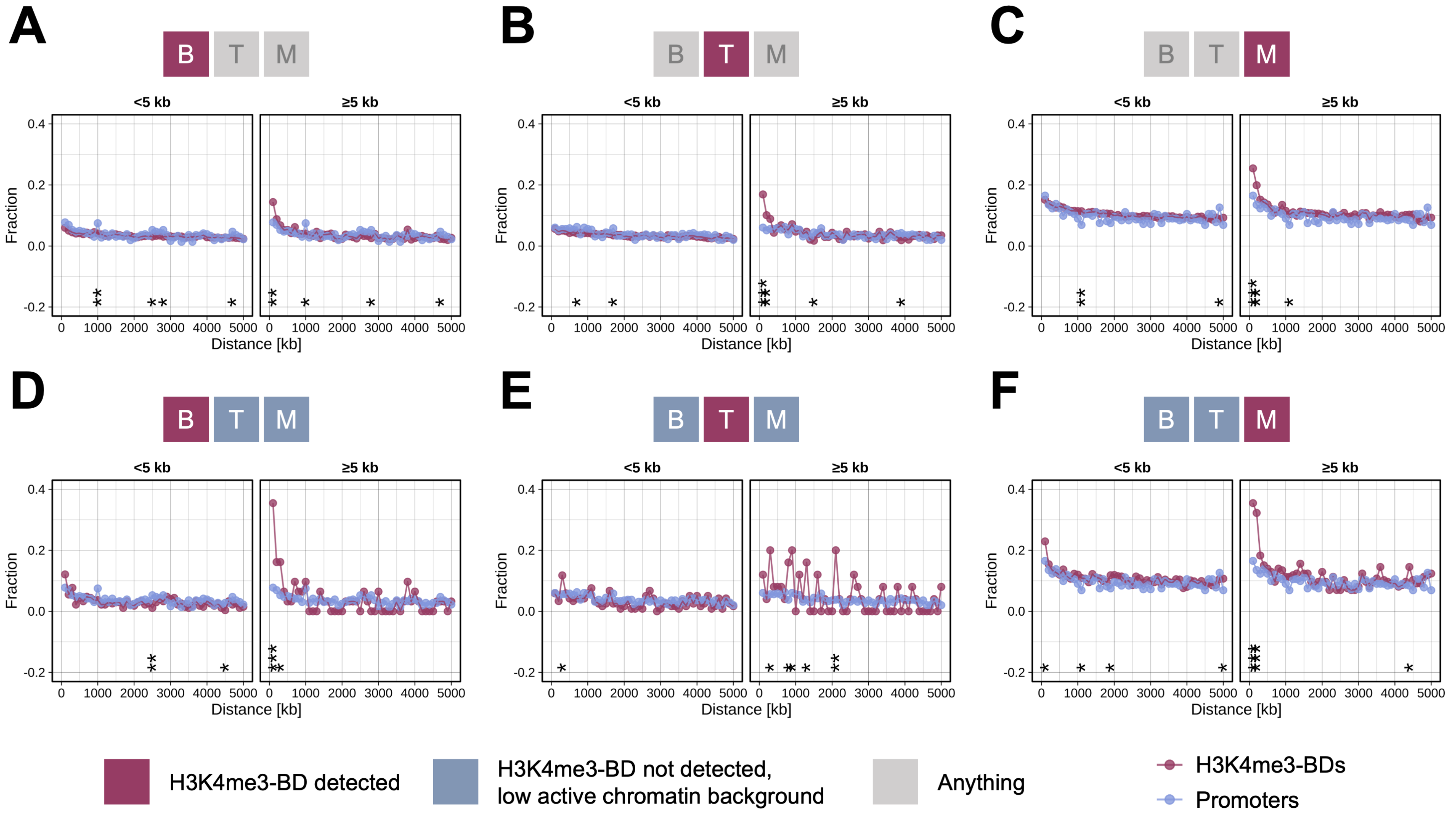
A
B
C
D
E
F
B T M
<5 kb
≥5 kb
Fraction
Distance [kb]
H3K4me3-BD detected
H3K4me3-BD not detected,
low active chromatin background
Anything
H3K4me3-BDs
Promoters

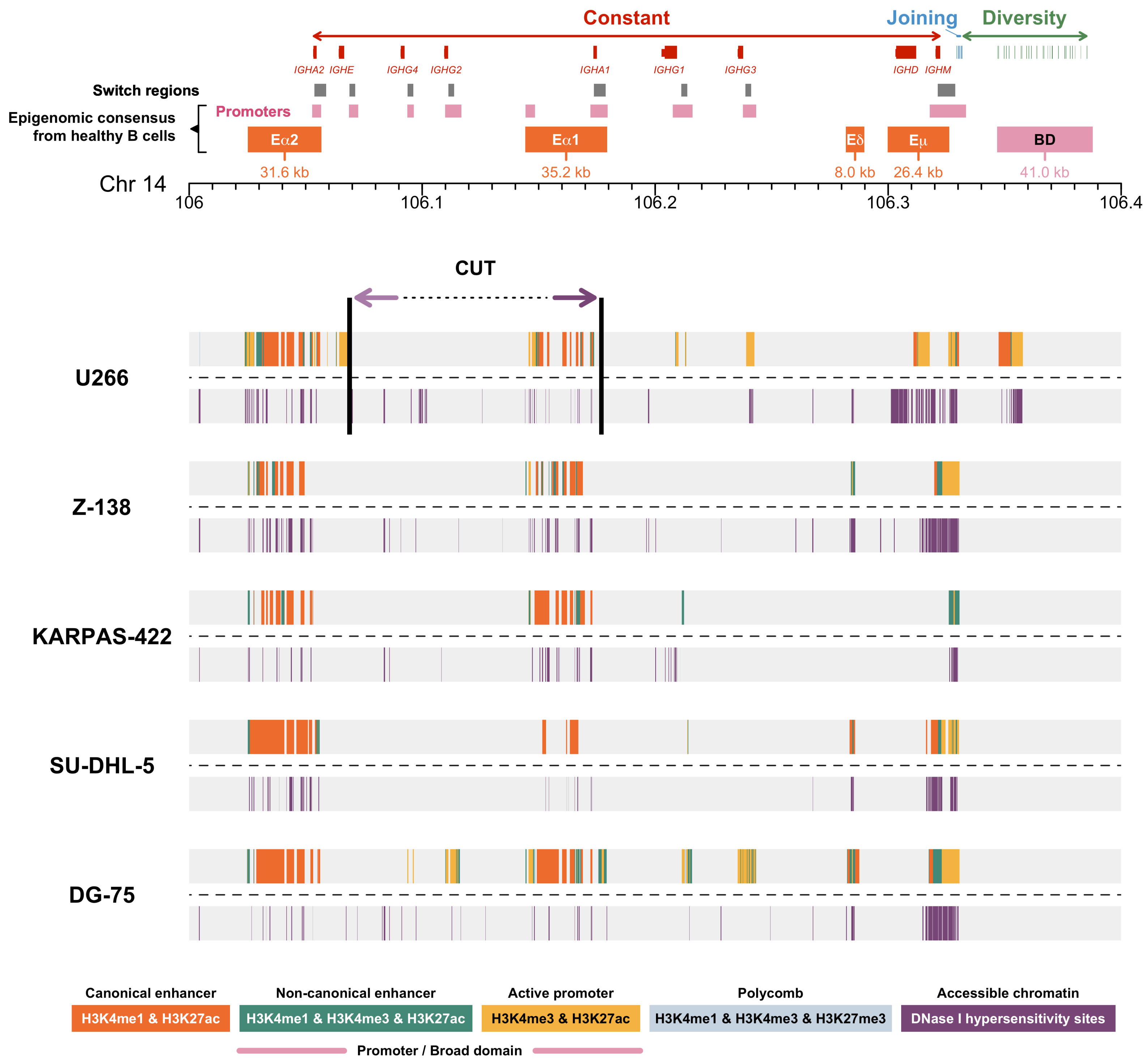

Constant
Joining
Diversity
IGHA2
IGHE
IGHG4
IGHG2
IGHA1
IGHG1
IGHG3
IGHD
IGHM
Switch regions
Epigenomic consensus from healthy B cells
Promoters
Eα2
Eα1
Eδ
Eμ
BD
31.6 kb
35.2 kb
8.0 kb
26.4 kb
41.0 kb
Chr 14
106
106.1
106.2
106.3
106.4
CUT
U266
Z-138
KARPAS-422
SU-DHL-5
DG-75
Canonical enhancer
H3K4me1 & H3K27ac
Non-canonical enhancer
H3K4me1 & H3K4me3 & H3K27ac
Active promoter
H3K4me3 & H3K27ac
Polycomb
H3K4me1 & H3K4me3 & H3K27me3
Accessible chromatin
DNase I hypersensitivity sites
Promoter / Broad domain

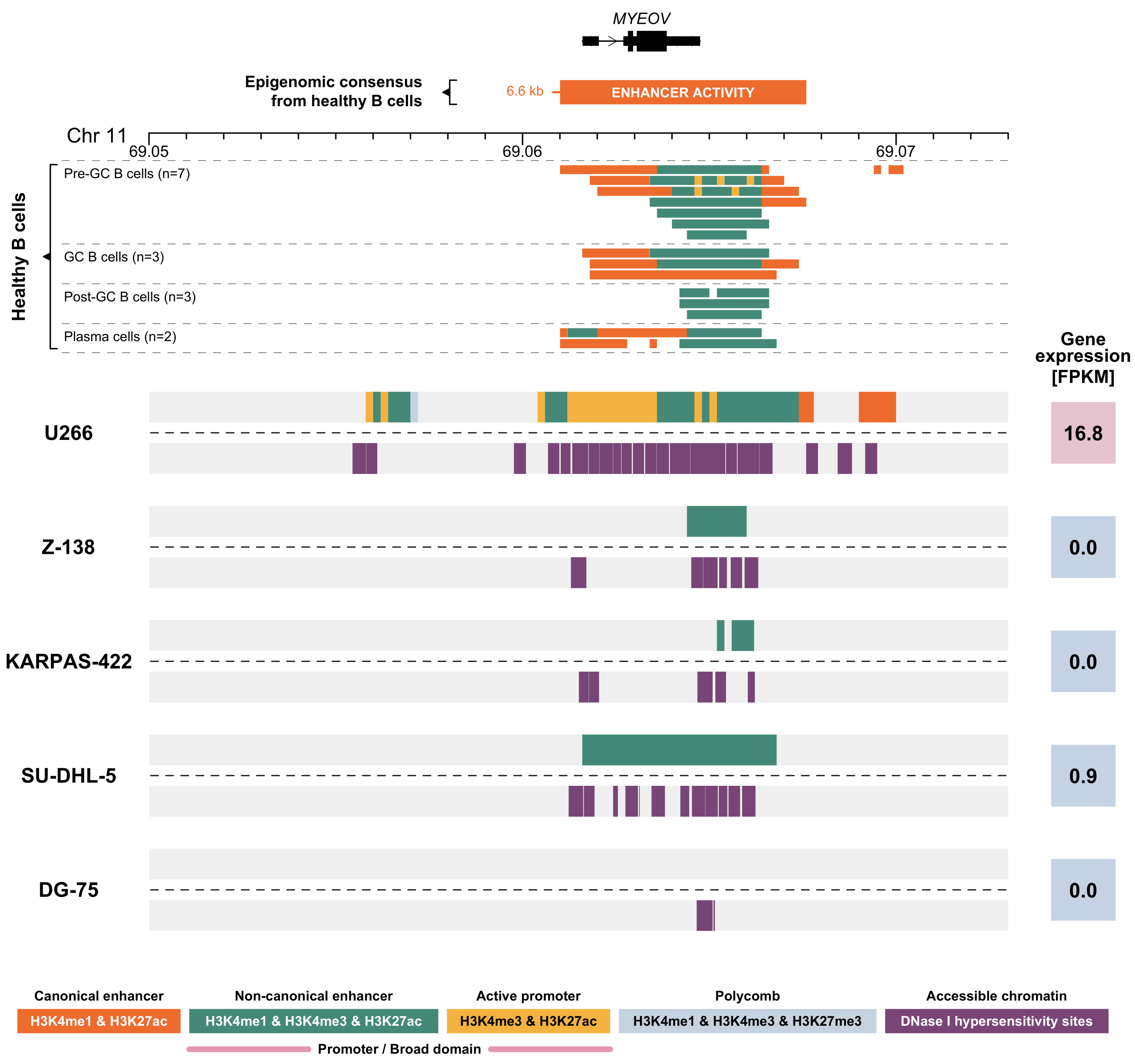

MYEOV
Epigenomic consensus from healthy B cells
6.6 kb
ENHANCER ACTIVITY
Chr 11
69.05
69.06
69.07
Healthy B cells
Pre-GC B cells (n=7)
GC B cells (n=3)
Post-GC B cells (n=3)
Plasma cells (n=2)
Gene expression [FPKM]
U266
16.8
Z-138
0.0
KARPAS-422
0.0
SU-DHL-5
0.9
DG-75
0.0
Canonical enhancer
H3K4me1 & H3K27ac
Non-canonical enhancer
H3K4me1 & H3K4me3 & H3K27ac
Active promoter
H3K4me3 & H3K27ac
Polycomb
H3K4me1 & H3K4me3 & H3K27me3
Accessible chromatin
DNase I hypersensitivity sites
Promoter / Broad domain

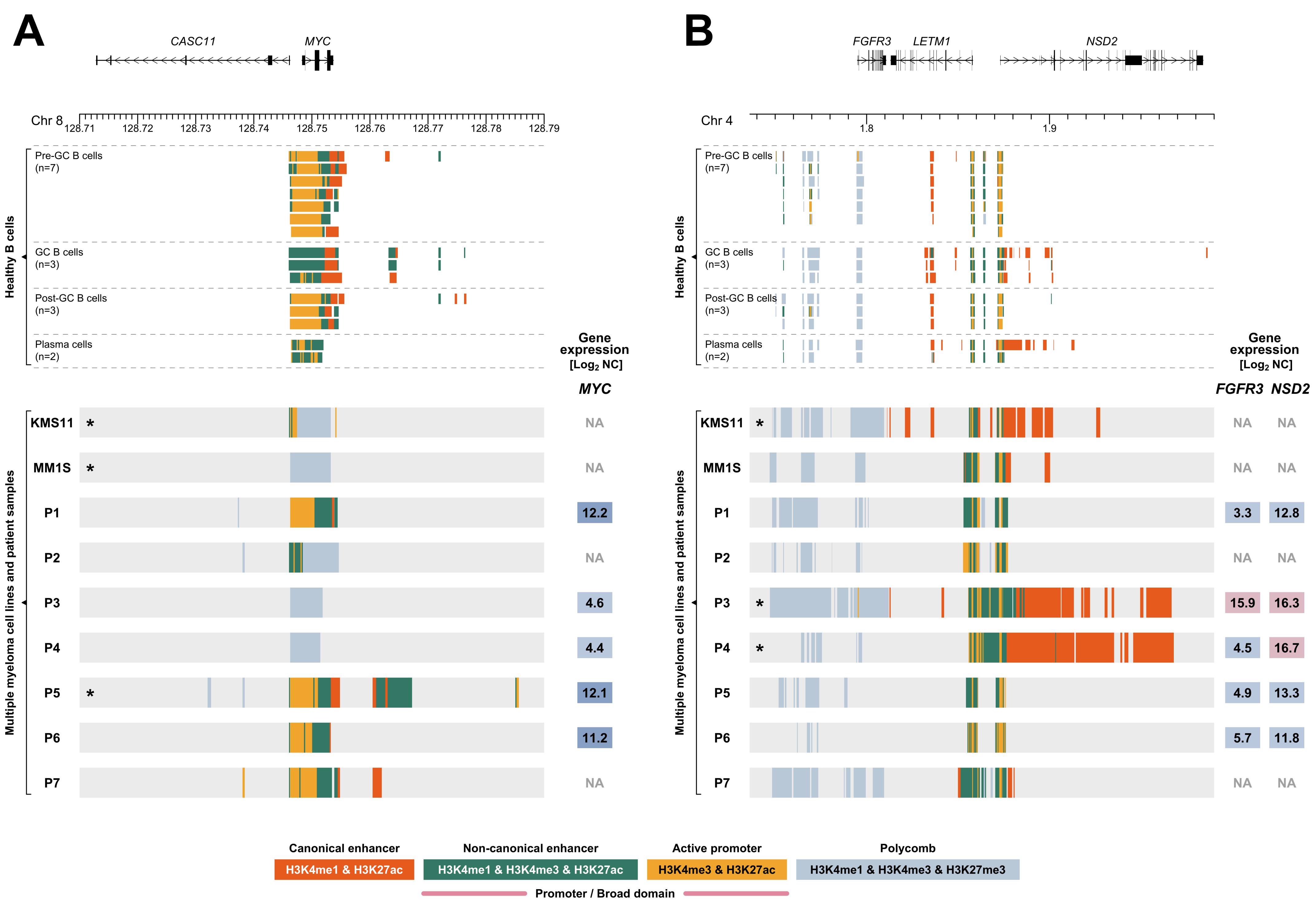

A
CASC11
MYC
Chr 8
128.71
128.72
128.73
128.74
128.75
128.76
128.77
128.78
128.79
Healthy B cells
Pre-GC B cells (n=7)
GC B cells (n=3)
Post-GC B cells (n=3)
Plasma cells (n=2)
Gene expression [Log2 NC]
MYC
Multiple myeloma cell lines and patient samples
KMS11 * NA
MM1S * NA
P1 12.2
P2 NA
P3 4.6
P4 4.4
P5 * 12.1
P6 11.2
P7 NA
B
FGFR3
LETM1
NSD2
Chr 4
1.8
1.9
Healthy B cells
Pre-GC B cells (n=7)
GC B cells (n=3)
Post-GC B cells (n=3)
Plasma cells (n=2)
Gene expression [Log2 NC]
FGFR3 NSD2
Multiple myeloma cell lines and patient samples
KMS11 * NA NA
MM1S NA NA
P1 3.3 12.8
P2 NA NA
P3 * 15.9 16.3
P4 * 4.5 16.7
P5 4.9 13.3
P6 5.7 11.8
P7 NA NA
Canonical enhancer
H3K4me1 & H3K27ac
Non-canonical enhancer
H3K4me1 & H3K4me3 & H3K27ac
Active promoter
H3K4me3 & H3K27ac
Polycomb
H3K4me1 & H3K4me3 & H3K27me3
Promoter / Broad domain

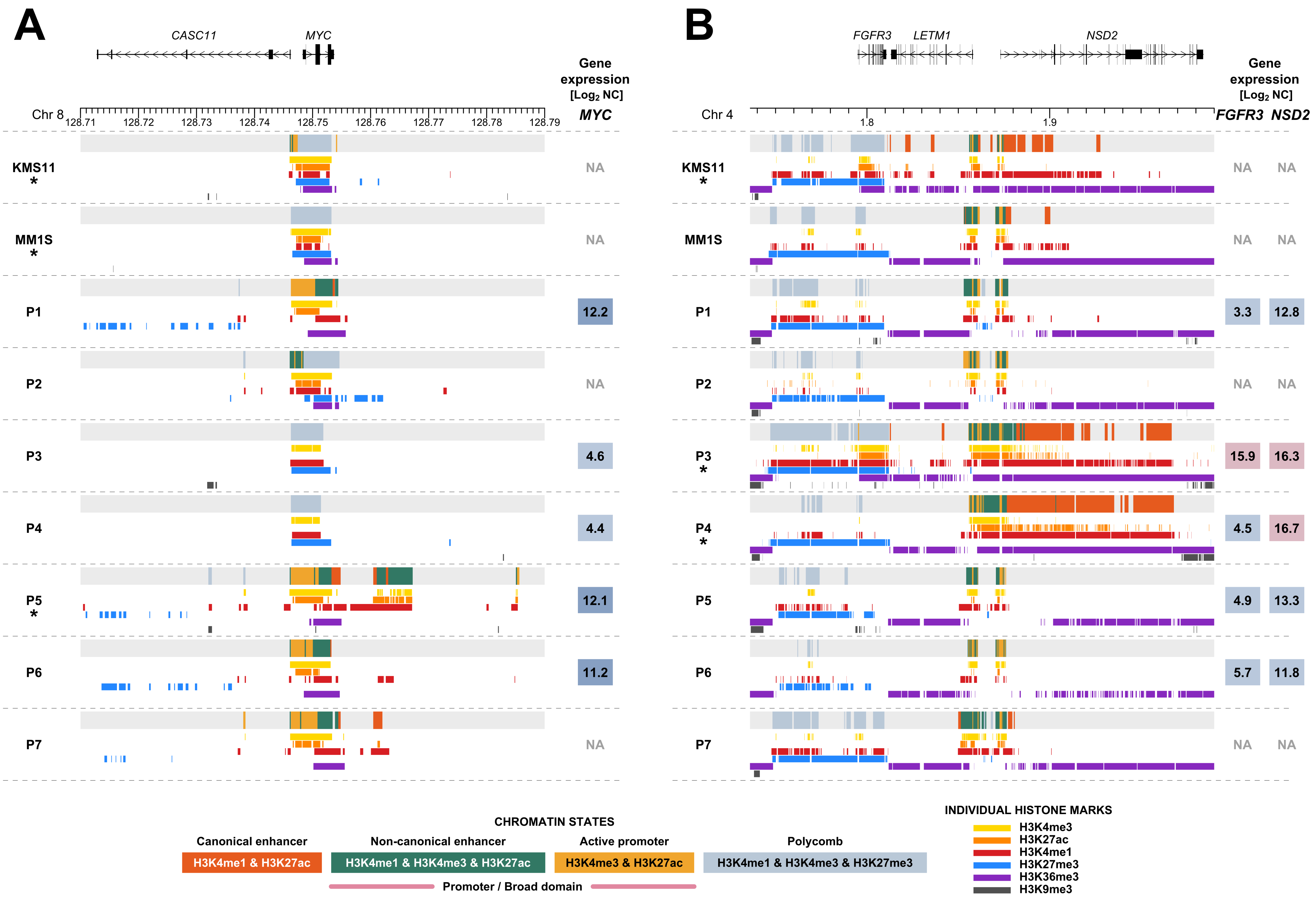
A
CASC11
MYC
Gene expression [Log2 NC]
MYC
Chr 8
128.71
128.72
128.73
128.74
128.75
128.76
128.77
128.78
128.79
KMS11 *
MM1S *
P1
P2
P3
P4
P5 *
P6
P7
NA
NA
12.2
NA
4.6
4.4
12.1
11.2
NA
B
FGFR3
LETM1
NSD2
Gene expression [Log2 NC]
Chr 4
1.8
1.9
FGFR3
NSD2
KMS11 *
MM1S
P1
P2
P3 *
P4 *
P5
P6
P7
NA NA
NA NA
3.3 12.8
NA NA
15.9 16.3
4.5 16.7
4.9 13.3
5.7 11.8
NA NA
CHROMATIN STATES
Canonical enhancer
H3K4me1 & H3K27ac
Non-canonical enhancer
H3K4me1 & H3K4me3 & H3K27ac
Active promoter
H3K4me3 & H3K27ac
Polycomb
H3K4me1 & H3K4me3 & H3K27me3
Promoter / Broad domain
INDIVIDUAL HISTONE MARKS
H3K4me3
H3K27ac
H3K4me1
H3K27me3
H3K36me3
H3K9me3

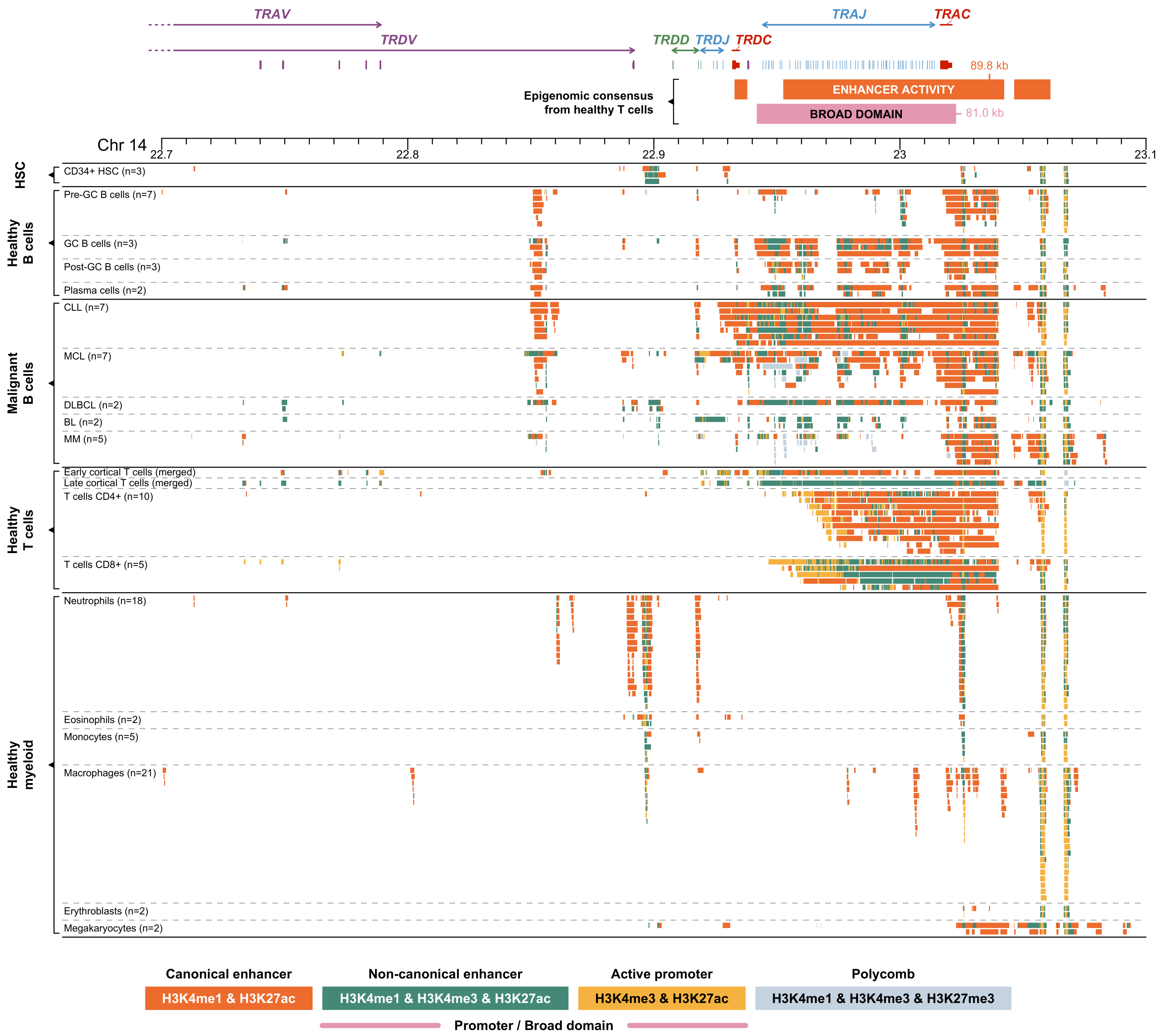

TRAV
TRDV
TRDD
TRDJ
TRDC
TRAJ
TRAC
89.8 kb
ENHANCER ACTIVITY
BROAD DOMAIN
81.0 kb
Epigenomic consensus from healthy T cells
Chr 14
22.7
22.8
22.9
23
23.1
HSC
CD34+ HSC (n=3)
Healthy B cells
Pre-GC B cells (n=7)
GC B cells (n=3)
Post-GC B cells (n=3)
Plasma cells (n=2)
Malignant B cells
CLL (n=7)
MCL (n=7)
DLBCL (n=2)
BL (n=2)
MM (n=5)
Healthy T cells
Early cortical T cells (merged)
Late cortical T cells (merged)
T cells CD4+ (n=10)
T cells CD8+ (n=5)
Healthy myeloid
Neutrophils (n=18)
Eosinophils (n=2)
Monocytes (n=5)
Macrophages (n=21)
Erythroblasts (n=2)
Megakaryocytes (n=2)
Canonical enhancer
H3K4me1 & H3K27ac
Non-canonical enhancer
H3K4me1 & H3K4me3 & H3K27ac
Active promoter
H3K4me3 & H3K27ac
Polycomb
H3K4me1 & H3K4me3 & H3K27me3
Promoter / Broad domain

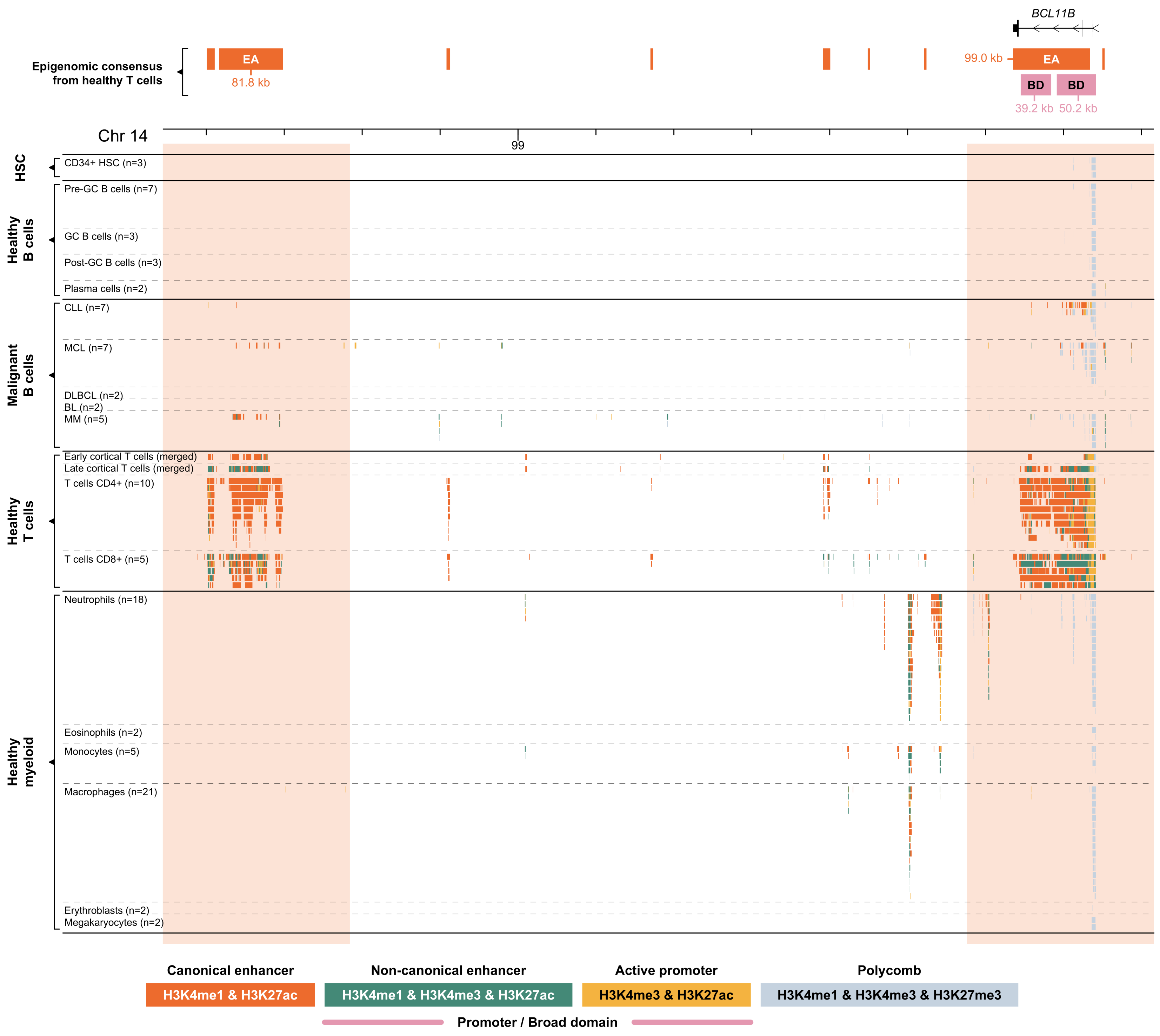

BCL11B
Epigenomic consensus from healthy T cells
EA
81.8 kb
99.0 kb
EA
BD
BD
39.2 kb
50.2 kb
Chr 14
99
HSC
CD34+ HSC (n=3)
Healthy B cells
Pre-GC B cells (n=7)
GC B cells (n=3)
Post-GC B cells (n=3)
Plasma cells (n=2)
Malignant B cells
CLL (n=7)
MCL (n=7)
DLBCL (n=2)
BL (n=2)
MM (n=5)
Healthy T cells
Early cortical T cells (merged)
Late cortical T cells (merged)
T cells CD4+ (n=10)
T cells CD8+ (n=5)
Healthy myeloid
Neutrophils (n=18)
Eosinophils (n=2)
Monocytes (n=5)
Macrophages (n=21)
Erythroblasts (n=2)
Megakaryocytes (n=2)
Canonical enhancer
H3K4me1 & H3K27ac
Non-canonical enhancer
H3K4me1 & H3K4me3 & H3K27ac
Active promoter
H3K4me3 & H3K27ac
Polycomb
H3K4me1 & H3K4me3 & H3K27me3
Promoter / Broad domain

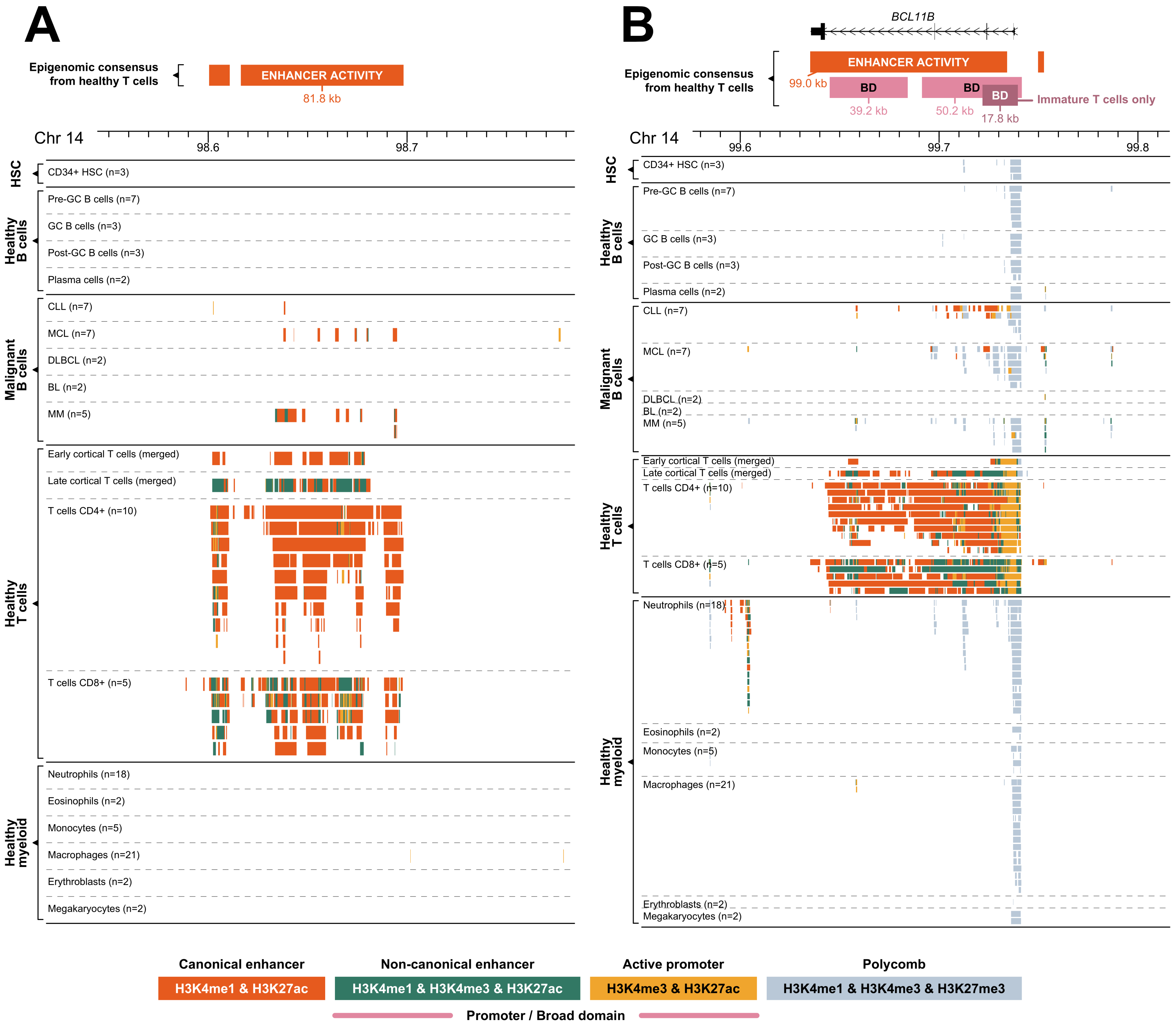

A
Epigenomic consensus from healthy T cells
ENHANCER ACTIVITY
81.8 kb
Chr 14
98.6
98.7
HSC
CD34+ HSC (n=3)
Healthy B cells
Pre-GC B cells (n=7)
GC B cells (n=3)
Post-GC B cells (n=3)
Plasma cells (n=2)
Malignant B cells
CLL (n=7)
MCL (n=7)
DLBCL (n=2)
BL (n=2)
MM (n=5)
Healthy T cells
Early cortical T cells (merged)
Late cortical T cells (merged)
T cells CD4+ (n=10)
T cells CD8+ (n=5)
Healthy myeloid
Neutrophils (n=18)
Eosinophils (n=2)
Monocytes (n=5)
Macrophages (n=21)
Erythroblasts (n=2)
Megakaryocytes (n=2)
B
BCL11B
Epigenomic consensus from healthy T cells
ENHANCER ACTIVITY
99.0 kb
BD
BD
BD
39.2 kb
50.2 kb
17.8 kb
Immature T cells only
Chr 14
99.6
99.7
99.8
HSC
CD34+ HSC (n=3)
Healthy B cells
Pre-GC B cells (n=7)
GC B cells (n=3)
Post-GC B cells (n=3)
Plasma cells (n=2)
Malignant B cells
CLL (n=7)
MCL (n=7)
DLBCL (n=2)
BL (n=2)
MM (n=5)
Healthy T cells
Early cortical T cells (merged)
Late cortical T cells (merged)
T cells CD4+ (n=10)
T cells CD8+ (n=5)
Healthy myeloid
Neutrophils (n=18)
Eosinophils (n=2)
Monocytes (n=5)
Macrophages (n=21)
Erythroblasts (n=2)
Megakaryocytes (n=2)
Canonical enhancer
H3K4me1 & H3K27ac
Non-canonical enhancer
H3K4me1 & H3K4me3 & H3K27ac
Active promoter
H3K4me3 & H3K27ac
Polycomb
H3K4me1 & H3K4me3 & H3K27me3
Promoter / Broad domain

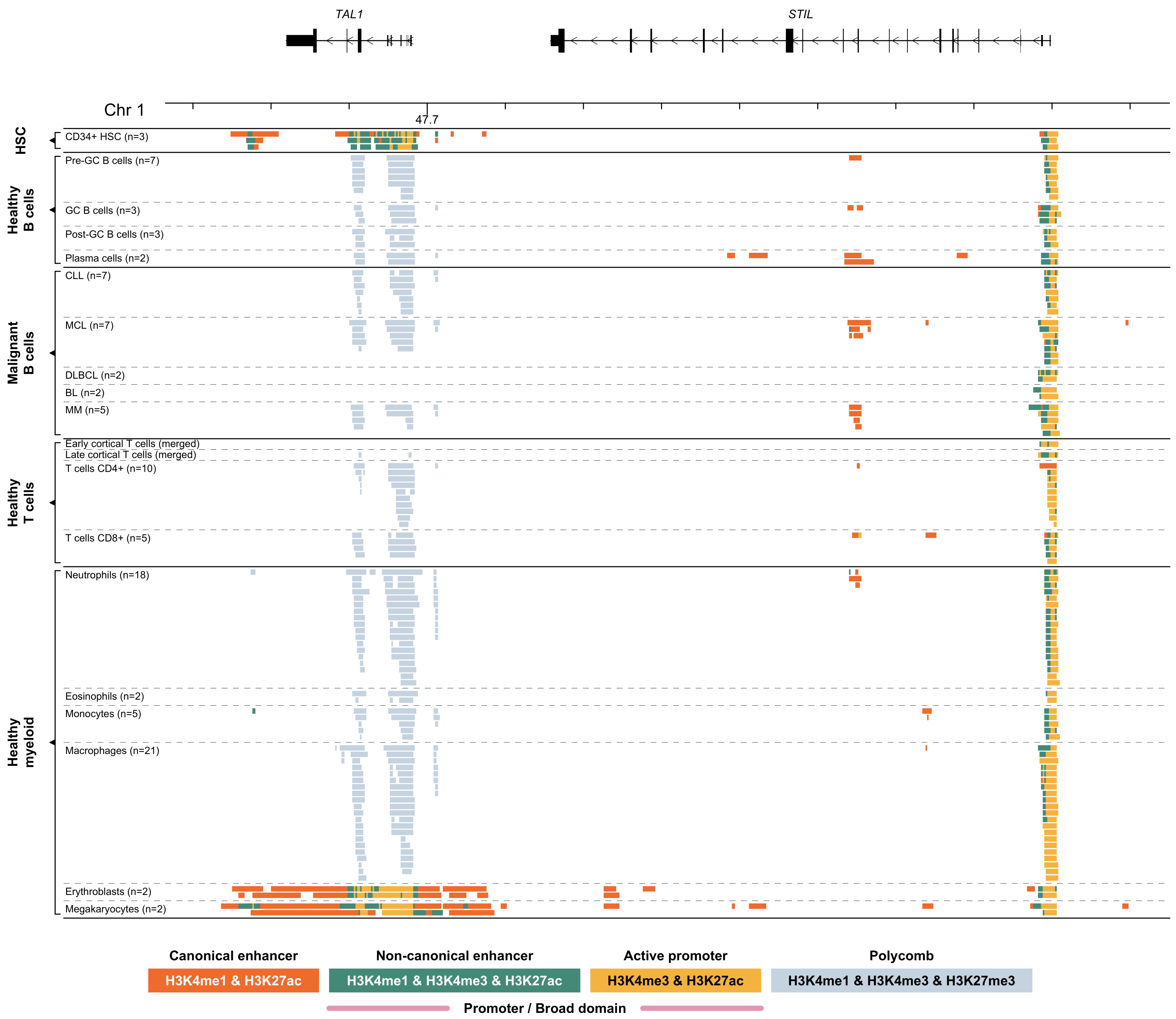

TAL1
STIL
Chr 1
47.7
HSC
CD34+ HSC (n=3)
Healthy B cells
Pre-GC B cells (n=7)
GC B cells (n=3)
Post-GC B cells (n=3)
Plasma cells (n=2)
Malignant B cells
CLL (n=7)
MCL (n=7)
DLBCL (n=2)
BL (n=2)
MM (n=5)
Healthy T cells
Early cortical T cells (merged)
Late cortical T cells (merged)
T cells CD4+ (n=10)
T cells CD8+ (n=5)
Healthy myeloid
Neutrophils (n=18)
Eosinophils (n=2)
Monocytes (n=5)
Macrophages (n=21)
Erythroblasts (n=2)
Megakaryocytes (n=2)
Canonical enhancer
H3K4me1 & H3K27ac
Non-canonical enhancer
H3K4me1 & H3K4me3 & H3K27ac
Active promoter
H3K4me3 & H3K27ac
Polycomb
H3K4me1 & H3K4me3 & H3K27me3
Promoter / Broad domain

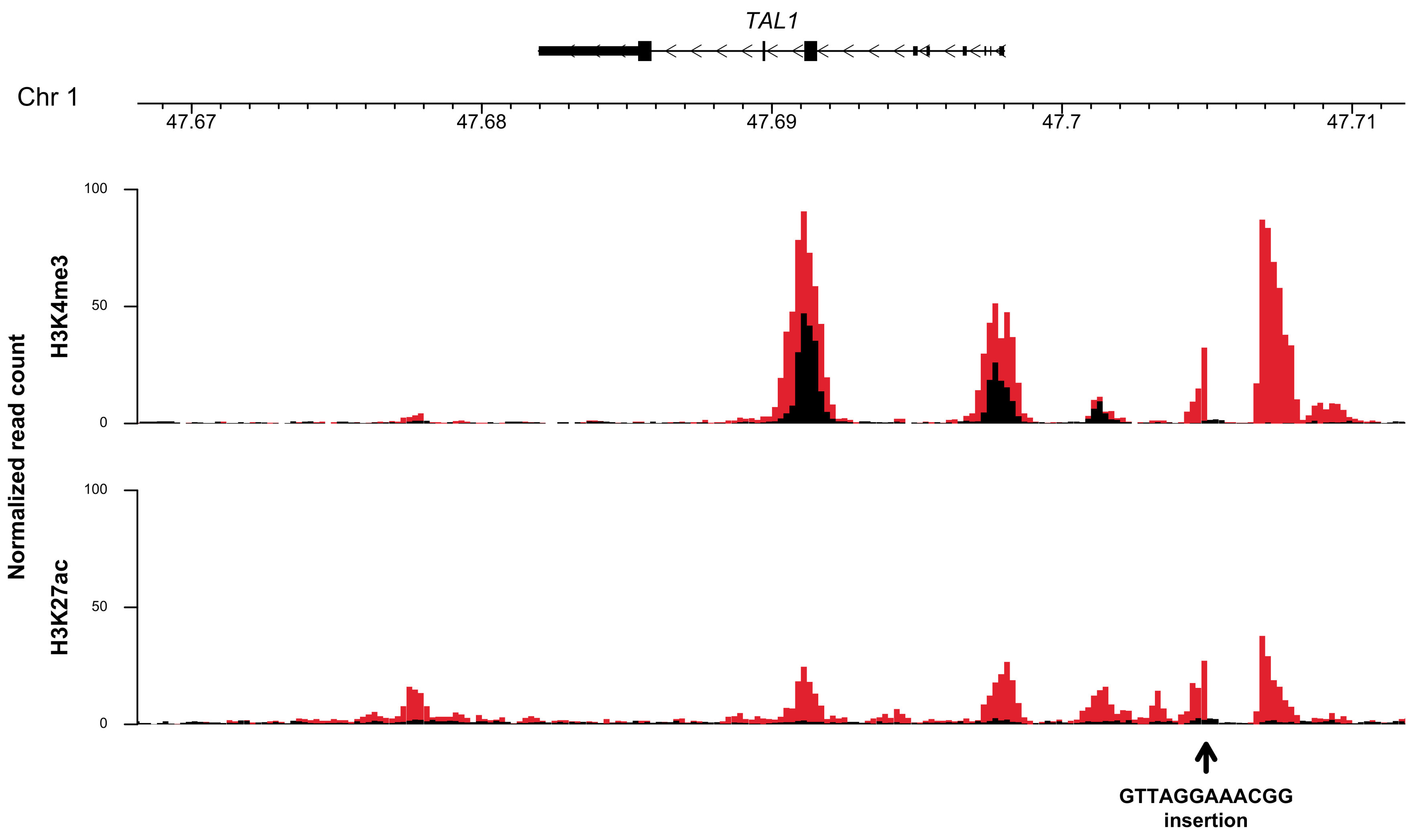
TAL1
Chr 1
47.67
47.68
47.69
47.7
47.71
Normalized read count
H3K4me3
H3K27ac
100
50
0
100
50
0
GTTAGGAAACGG
insertion
